# Antagonistic insulin mimetics lock the insulin receptor in an alternative apo-state

**DOI:** 10.1101/2025.08.05.667158

**Authors:** Martin Polák, Irena Selicharová, Matyáš Pinkas, María Soledad Garre Hernández, Benjamin Fabre, Marta Lubos, Lenka Žáková, Michal Grzybek, Ingmar B. Schäfer, Ünal Coskun, Jiří Jiráček, Jiří Nováček

**Author notes:** Correspondence to: Jiří Nováček and Jiří Jiráček.

## Abstract

Despite significant advancements in high-resolution structural analysis of activated human insulin receptor (IR), the molecular mechanisms underlying its conformational plasticity that govern the transition from the *apo* state to the activated state are still not well understood. This leaves critical aspects of IR regulation unclear. Here, we reveal the mechanism by which the insulin mimetics Ada, Trim, and S661 fully inhibit the insulin receptor. The receptor is stabilized in a yet structurally un–described ∩–shaped conformation which is induced by antagonist binding between the L1 and FnIII-1’ domains. In contrast to insulin-bound IR structures, the α-CT helix is not observable in the ∩ conformation, and the membrane-proximal regions of the FnIII-3 domains are >10 nm apart, which prohibits transmembrane signal transduction and kinase domain activation. Analysis of apo-IR electron cryo-microscopy data indicates that the ∩-shaped state is one of several metastable apo-IR conformations. These findings underscore the intrinsic conformational dynamics of apo-IR and its role in integrating insulin binding and receptor activation.

## Introduction

Receptor tyrosine kinases (RTKs) are a family of pivotal membrane receptors that regulate various cellular processes, including growth, differentiation, and metabolism. RTK signal transduction is often initiated at the cell surface by ligand-induced receptor dimerization, which results in conformational changes and the subsequent activation of downstream signaling pathways (1). The insulin receptor (IR), the insulin-like growth factor receptor (IGF-1R), and the insulin receptor-related receptor (IRR) form a unique RTK subgroup (2, 3). These RTKs are delivered to the plasma membrane as disulfide-bonded, preformed dimers in their unliganded apo-structure. This necessitates an autoinhibitory mechanism to prevent ligand-independent activation of signaling pathways, but the details of this mechanism remain incompletely understood. The functional IR is a key regulator of animal physiology (4). IR consists of two protomers, each of which has one extracellular α-subunit and one trans-membrane β-subunit. These subunits are linked by disulfide bonds to form a heterotetrameric complex (αα’ββ’). The IR extracellular region (referred to as the ectodomain or IR-ECD) consists of the leucine-rich domain 1 (L1), the cysteine-rich domain (CR), the leucine-rich domain 2 (L2), and three fibronectin type III domains (FnIII-1, FnIII-2, and FnIII-3). The α-CT peptide, a long helical segment that stabilizes the IR structure, is a key structural element within the α subunit’s C-terminus, which plays a pivotal role in insulin binding and receptor activation. The β subunit contains the membrane-proximal FnIII-2 and FnIII-3 domains of the IR-ECD, followed by the IR transmembrane domain (IR-TMD), the intracellular juxtamembrane domain (JM), the tyrosine kinase domain (TKD), and the C-terminal tail (5). In mammals, the *INSR* gene produces two IR isoforms, IR-A and IR-B, through alternative splicing (6).

Structural studies have revealed sequential, ligand-induced rearrangements during IR activation. Current consensus from cryo-EM and X-ray crystallography studies of IR-ECD and full-length IR in detergent micelles suggests a continuum of conformational states dependent on insulin occupancy (7-14). IR-ECD conformers associated with ligand-induced activation begin with the inactive, Λ-shaped apo structure. Upon binding to one insulin, the asymmetric Γ-shaped structure is formed. This is followed by the asymmetric 𝒯-shaped structure with two to three bound insulins (13). Finally, the structure becomes the fully symmetric, ligand-saturated, T-shaped structure with four insulins (11, 12). Insulin binding and IR activation are mediated via two distinct receptor-binding sites on each protomer that engage differently in the conformers. Site-1 is located on the L1 domain and comprises the membrane-distal part of the FnIII-1′ domain and the α-CT′ helix. Site-2 is situated on the membrane-proximal region of the FnIII-1′ domain. It is generally assumed that synergistic engagement of both sites is essential for activation (15). Site-2 is thought to initially interact with insulin, positioning it for subsequent translocation to Site-1 (13). Thus, insulin binding destabilizes the auto-inhibited apo-IR structure, triggering sequential structural rearrangements of the active receptor that ultimately bring the transmembrane domains closer together, enabling auto-phosphorylation of the kinase domains. However, the structural dynamics of apo-IR, including its transition toward ligand-binding-competent states, such as the asymmetric, Γ-shaped conformation, remain poorly understood. Notably, both full-length IR in lipid nanodiscs and the isolated IR-ECD adopt conformations that differ from the canonical Λ-shaped structure in the apo state.

Phage display screens have identified insulin-non-related peptides that bind to IR with high affinity and modulate receptor signaling, either by activating or inhibiting it (16, 17). For instance, peptides such as S597 and S519 have been demonstrated to exhibit agonistic properties (16), while the S661 peptide has been shown to act as a potent antagonist (18). These peptides contain specific structural motifs that enable them to compete with endogenous α-CT peptides (which are part of Site-1 of the receptor) for binding to the L1 domain. They also contain motifs that bind to Site-2 of the receptor in the FnIII-1’ domain (see Table S1). We have previously developed bi- and tri-podal peptidomimetic compounds to target the IR. These compounds were synthesized by attaching distinct peptide sequences derived from the S597 and the S661 peptides to a central molecular scaffold. Using trimesic acid (19) and adamantane (20) as scaffold molecules, we have prepared Trim and Ada insulin mimetics, respectively. These compounds contain two key peptide sequences which bind to Site-1 and to Site-2. Both sequences are attached to the central scaffolds via their C-termini (Figure 1A, Table S1). Ada and Trim have been shown to bind to IR, yet they do not activate the receptor. Rather, they function as full IR antagonists.

**Figure 1.**
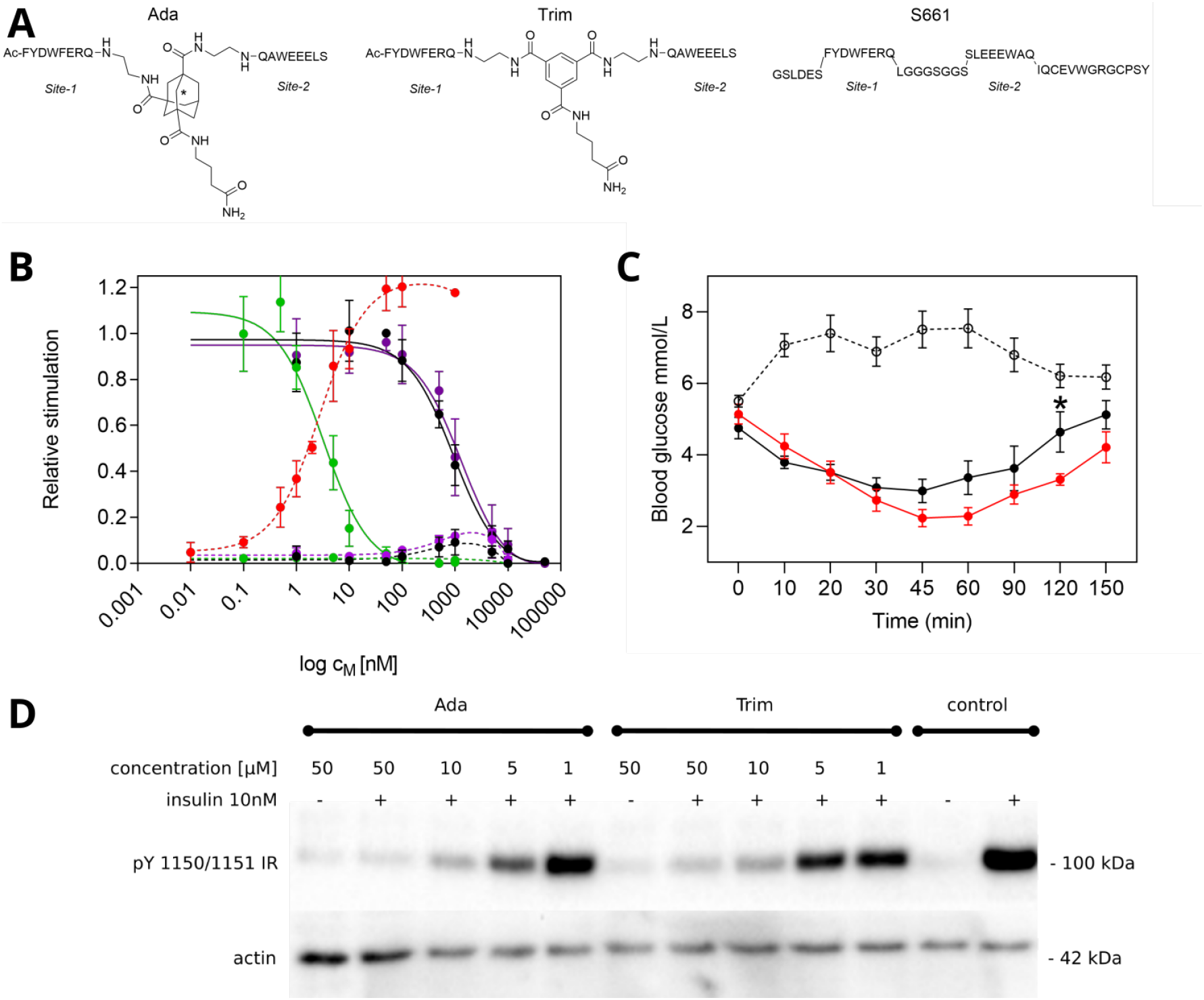
Schematic representation of the Ada, Trim, and S661 insulin mimetics (A). Ada is a mixture of two diastereoisomers (depicted by an asterisk). The minimal Site-1 (Ac-FYDWFERQ) and Site-2 (SLEEEWAQ) peptides are connected to the scaffold molecule via their C-termini in case of Ada and Trim. Activation of IR autophosphorylation by insulin (B, red dotted line) and inhibition of insulin-induced (10 nM) IR activation by Ada (B, black), Trim (B, violet), and S661 (B, green). Dotted lines show no effect of Ada, Trim, and S661 on the IR autophosphorylation in the absence of insulin. The results (± S.E.) of the insulin tolerance test (ITT) in male C57BL/6J mice (n=10 per group) is shown in C. Mice were fasted for 18h and then injected with either saline (dotted black line), human insulin at 0.75 U/kg (red line), or a combination of human insulin (0.75 U/kg) and Ada (125 U/kg) (black line). The black asterisk highlights difference between effect of human insulin and human insulin in presence of Ada (p = 0.0345, Multiple T-test). Representative Western blots for the abilities of Ada and Trim compounds to inhibit insulin stimulated IR-A phosphorylation (D). Cells were stimulated with 50 mM compounds alone (−) or with 50 - 1 mM compounds in the presence of 10 nM insulin (+) for 10 min. Membranes were cut at 75 kDa and 50 kDa standards and respective parts were developed with anti-phospho-IGF-1Rβ (Tyr1135/1136)/IRβ (Tyr1150/1151) antibody (Mr above 75 kDa) and with anti-actin antibody (Mr bellow 50 kDa).

In this study, we employed single particle electron cryo-microscopy (cryo-EM) to elucidate the mechanism of IR inhibition by Ada, Trim, and S661. The three antagonists induce an IR arrangement that has not yet been structurally characterized. This IR configuration is denoted ∩ conformation and it exhibits a high degree of similarity with 2D class averages observed for apo-IR earlier (11,21). Our cryo-EM data demonstrate that the helical segments of Ada, Trim, and S661 stabilize the IR L1 and FnIII-1’ domains in close proximity. Furthermore, the antagonist displaces the α-CT helix and the FnIII-2 domain from the L1 domain. This results in receptor stabilization in the ∩ conformation, thereby locking it in an inactive state. A thorough examination of apo-IR single particle cryo-EM data has revealed that, in addition to the previously identified Λ-shaped structure, a substantial proportion of the apo-IR conformational ensemble consists of other conformations that bear a striking resemblance to the Ada/Trim/S661-bound ∩-IR. In summary, the ∩ conformation represents a substantial reorganization of the apo-receptor in comparison to the Λ conformation. It involves the stabilization of Site-1 and Site-2 in close proximity, while maintaining a separation of more than 10 nm between the membrane proximal FnIII domains. This conformation manifests as an autoinhibited form, poised to engage ligands with a mixed site 1/2, analogous to the configuration described in the 𝒯-shaped structure (11, 13). Consequently, our findings offer not only a comprehensive structural framework for the full inhibition of IR by Ada, Trim, and S661, but also expand our understanding of apo-IR conformational dynamics and the regulation of activation.

## Results

### Insulin mimetics bind but do not activate IR and fully inhibit insulin-induced IR activation

We first performed *in vitro* binding assays which demonstrate that Ada and Trim bind to IR-A with dissociation constants (*K*_d_) of 395 nM and 1257 nM, respectively. In contrast, S661 binds with markedly higher affinity (*K*_d_ of approximately 2.6 nM; Tab. S1, Fig. S1). Ada, Trim and S661 have been shown to inhibit insulin-induced IR activation in a concentration-dependent manner. The competitive binding curves show curvilinear Scatchard plots (Fig. S1). These results are consistent with the presence of two binding sites. Furthermore, peptidomimetics Ada and Trim, akin to the S661 peptide, diminish insulin-stimulated IR phosphorylation to levels that are indistinguishable from the background in mouse embryonic fibroblasts (Fig 1B,D, Fig. S2). These findings corroborate the hypothesis that both Ada and Trim function as full IR antagonists, a property exhibited by S661. To assess the antagonistic effects *in vivo*, an insulin tolerance test was performed with Ada in mice (Fig. 1C). Mice injected with insulin demonstrated a substantial decrease in blood glucose levels. However, when insulin was co-administered with Ada, higher blood glucose levels were retained even after 45 minutes, indicating that Ada effectively attenuates insulin action *in vivo*.

### Apo-IR ectodomain exhibits extensive conformational dynamics

We then set out to investigate the molecular mechanism of IR inhibition by Ada, Trim, and S661. The single particle cryo-EM data was collected on both the inhibitor-bound and unliganded IR using the human IR-A ECD construct (residues 1-917, Fig. 2A), which was purified as previously described (11). The apo-IR samples exhibited well-dispersed particles, yet demonstrated substantial flexibility, as evidenced by the heterogeneity observed during 2D classification (Fig. 2B,C). Following a series of reference-free 2D classifications in combination with 3D classifications utilizing an *ab initio* routine in CryoSparc (22), a map corresponding to the previously described Λ-shaped IR structure was refined to 6 Å resolution (Fig. 2D, Fig. S3). The resulting map was interpreted through flexible fitting of the previously determined apo-IR structure stabilized by Site-1 deficient insulin (PDB: 7SL1, 15). The Λ-shape structure is stabilized by an extensive interaction interface between the FnIII-2’ and the L1+CR domains, as well as the α-CT helices, which bind to Site-1 on the L1 domain (Fig. 2D). Despite these stabilizing interactions, the apo-IR-A-ECD exhibited a high degree of intrinsic flexibility, which constrained the resolution of the map and refinement to higher resolution. Furthemore, only approximately 34.000 of the more than 1 million particles that passed the initial classification rounds contributed to the final refinement of Λ-shape structure. Interestingly, a considerable subset of the remaining particles still contributed to well-defined 2D class averages during the 2D classification rounds (Fig. 2C). However, these 2D class averages were clearly distinct from the Λ-shaped IR structure. This observation lends further support to the previous hypothesis that the Λ-shaped IR structure is not the sole meta-stable structural arrangement present in the apo-IR sample (11, 21). However, attempts to refine these non-Λ particles into high-resolution 3D models resulted only in low-resolution reconstructions (Fig. 2C, Fig. S3A), which could not be further interpreted unambiguously using molecular models. A 3D interpretation of these non-Λ conformational states became possible through the structures of the IR stabilized by Ada, Trim, and S661 (*vide infra*), providing first insights into the apo-IR conformations at equilibrium.

**Figure 2.**
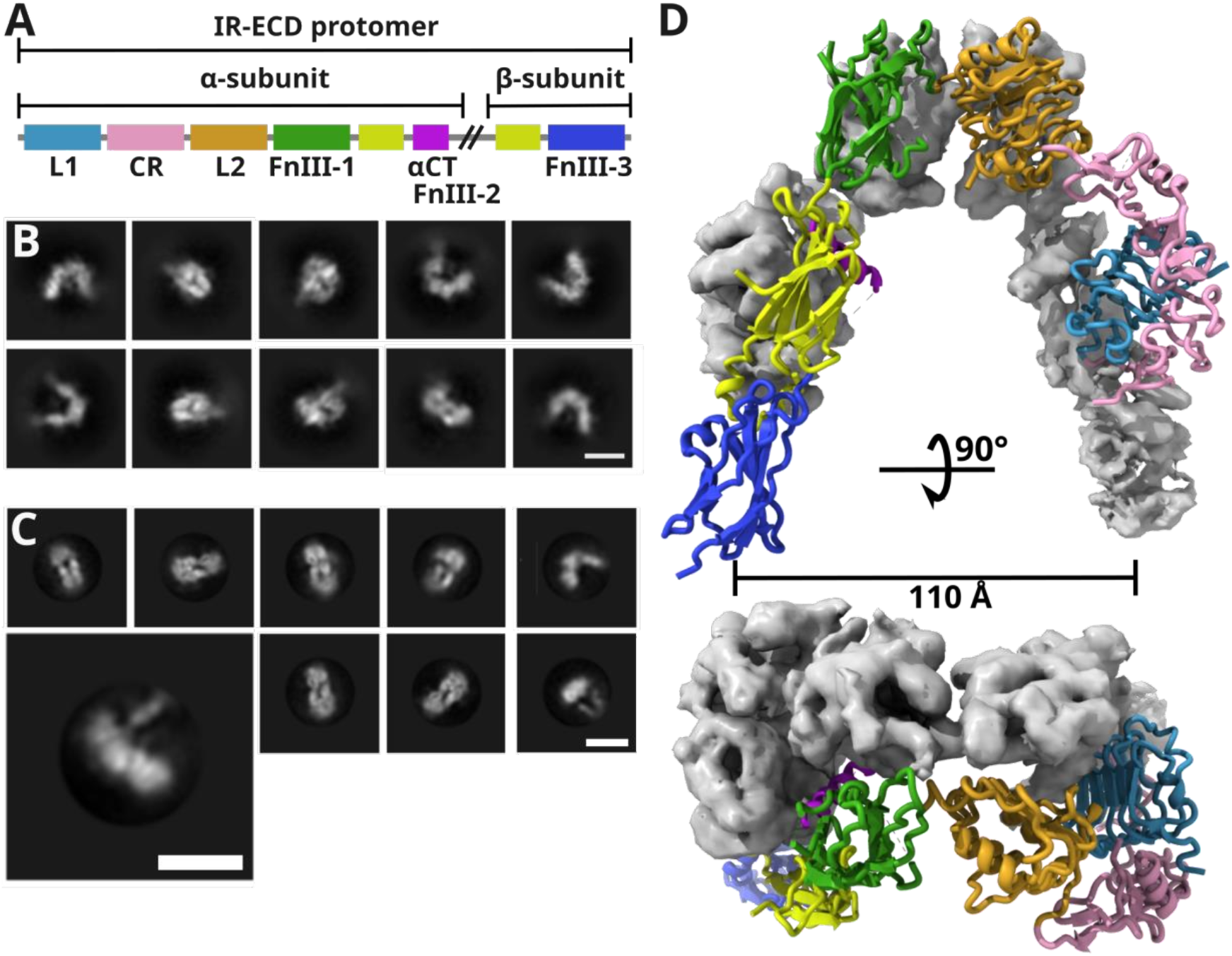
The domain composition of the IR-ECD construct used in this study (A). The color coding of individual domains is used throughout the manuscript to highlight individual domains in the models and cryo-EM maps (L1 domain in light-blue, CR domain in pink, L2 domain in orange, FnIII-1 in green, FnIII-2 in yellow, FnIII-3 in blue). Selected reference-free 2D class averages corresponding to the Λ-shaped apo-IR conformation (B). Well-defined 2D class-averages observed in the apo-IR data corresponding to other conformation than Λ-shaped apo-IR (C). The highlighted class-average image represents a structure, which was observed for antagonist bound IR-ECD under unsaturated ligand conditions. Scale bars in B and C correspond to 10 nm. Side (top) and top (bottom) view of the Λ-shaped apo-IR-ECD cryo-EM map (D) refined to 6 Å resolution and interpreted by flexible fitting of the apo-IR structure stabilized by Site-1 deficient insulin (PDB: 7SL1, 15). A map is shown for one protomer, whereas the fitted model is shown for the latter protomer. Color coding corresponds to the domain assignment shown in panel A.

### Antagonist binding stabilizes IR in ∩-shaped structure

In order to investigate the structural effects of antagonistic binding, IR-A-ECD (1.1 µM was the most suitable concentration for recording single particle micrographs in our hands) was mixed with Ada, or Trim in 100-fold molar excess prior to single particle cryo-EM analysis. The IR-ECD:Ada complex (Fig. 3A, Fig. S4) yielded a cryo-EM map with an average resolution of 3.0 Å (FSC0.143), while the ECD:Trim complex was resolved to 3.2 Å resolution (FSC0.143, Fig. S5). A close examination of the maps revealed that both Ada and Trim binding resulted in highly analogous structural organization (Fig. 4A,B). To streamline the subsequent discussion, the term “antagonist” will be used to collectively refer to these two compounds. As is typical of RTKs, IR-ECD is known to be significantly glycosylated and our cryo-EM maps have revealed additional densities at 10 asparagine residues, which we attribute to glycosylation (Fig. S6).

**Figure 3.**
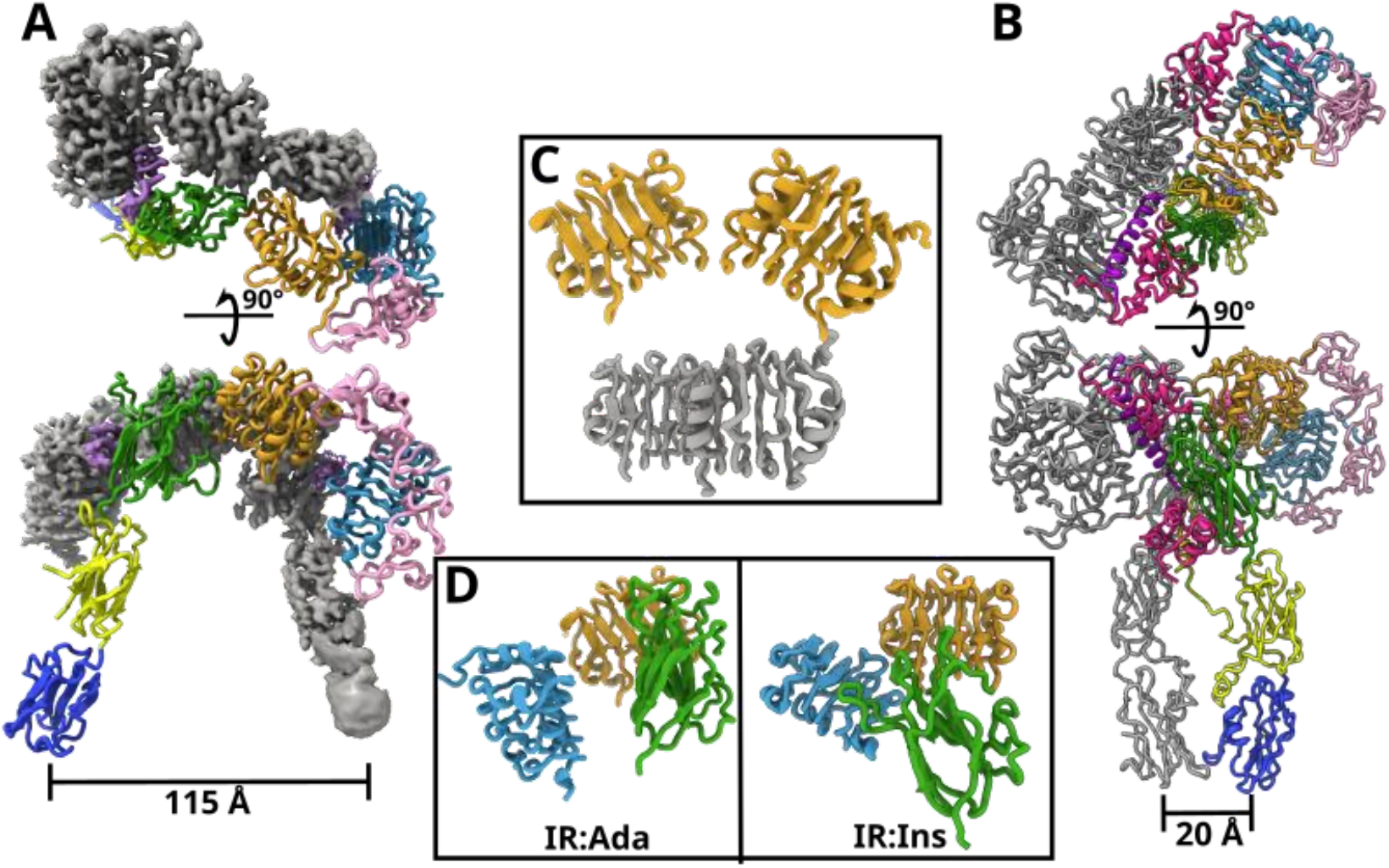
Cryo-EM structure of the IR-ECD:Ada complex in the ∩ conformation depicted in the mixed model/map visualisation (A). The Ada peptidomimetics is highlighted in violet. Insulin activated T-shaped IR-ECD (PDB: 6SOF) with insulin molecules highlighted in pink (B). The β-barrels of the L2 domains are parallel in the insulin-bound structure (in grey) whereas they are rotated by 70° in the IR-ECD:Ada (in orange) (C). In contrast to insulin-bound structure, FnIII-1 domain is positioned in similar distance with respect to the membrane in upon IR-ECD:Ada binding (D). The individual IR domains are color-coded in accordance with the scheme in Fig. 2A. Domains from different IR protomers are not distinguished in panels C and D.

**Figure 4.**
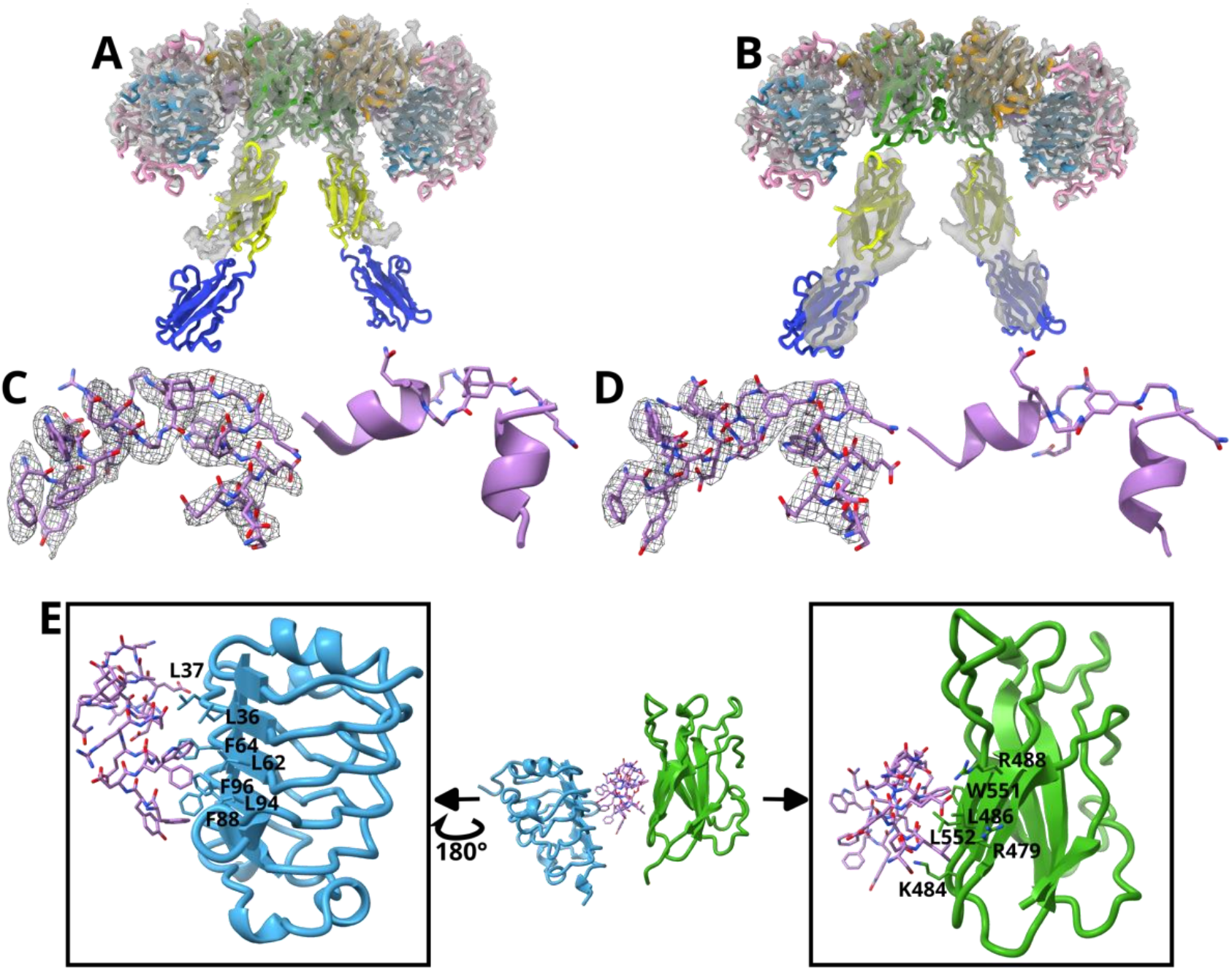
IR-ECD:Ada (A) and IR-ECD:Trim (B) (color coding corresponds to Fig. 2A,D in both cases; domains from different IR protomers are not distinguished), rigid regions comprising L1, CR, L2, and FnIII-1 domains for both protomers were refined to 3.0 Å and 3.2 Å resolution, respectively. The FnIII-2/3 domains retained higher flexibility which prevented refinement of these regions to high resolution. Instead, FnIII-2/3 domains were interpreted by flexible fitting of their previously determined structures (PDB: 6SOF, 11) into cryo-EM map. Ada (C) and Trim (D) are composed of two helical motifs connected through their C-termini into a V-shaped structure by adamantane or trimesic acid scaffolds. The Site-1 helix (composed of N’-FYDWFERQ-C’ amino acids) binds to the hydrophobic pocket on L1 domain (E). In addition, the Site-2 helix (N’-SLEEEWAQ-C’) binds to hydrophobic and positively charged region on FnIII-1 domain (E).

The antagonist-bound IR-ECD forms a pseudo-symmetric complex with two antagonist molecules bound to a single IR dimer, thereby stabilizing the receptor in a distinct conformation, which we designate as the ∩ conformation (Fig. 3A). In this configuration, the L1, CR, L2, and FnIII-1 domains form a compact symmetric head in the membrane-distal region. In contrast, the FnIII-2 and FnIII-3 domains exhibit asymmetry and a notable degree of flexibility. For the membrane-proximal FnIII-2/3 domain densities, the cryo-EM densities were insufficient for modelling; these regions were instead interpreted by flexible fitting of the previously solved structure of insulin bound IR-ECD (PDB: 6SOF). Notably, the densities corresponding to the α-CT helices, which typically stabilize interactions between the L1 and L2 domains in the insulin-bound structures, were entirely absent in the antagonist-bound maps.

Overall, the structure of the IR-ECD:antagonist complex (Fig. 3A) is in a more opened conformation compared to the insulin-bound IR-ECD (Fig. 3B). The receptor “head”, which consists of the L1, CR, L2, and FnIII-1 domains, forms an elongated structure measuring ∼145 Å in length and ∼60 Å in width. In contrast to insulin-bound IR-ECD, the binding of the antagonist causes the β-barrel of the L1 domain to adopt an orientation that is 70° tilted to the anticipated membrane plane (compared to 30° in insulin-bound IR-ECD, Fig. 3D). This distinct orientation is likely attributable to two factors: the stabilization of L1 by its interaction with the FnIII-1 domain mediated by the antagonist molecule, and the absence of stabilizing interactions with the α-CT helix, which is displaced upon antagonist binding. Additionally, the β-barrels of the L2 domains do not align parallel to the membrane as observed in the insulin-bound IR-ECD (Fig. 3C). In contrast, the N-termini of the latter are rotated ∼34° away from the membrane (Fig. 3C). Furthermore, the FnIII-1 becomes more integrated into the receptor head upon binding of the antagonist (Fig. 3D). Collectively, these structural rearrangements result in a receptor head that is flatter and wider than that of the insulin-bound IR (Fig. 3A,B). In this configuration, the L1, CR, L2, and FnIII-1’ form a ring-like structure. It is noteworthy that the L1, CR, L2 and FnIII-1’ domains are arranged in one plane, which is tilted approximately 15° relative to the plane formed by their counterparts, the L1’, CR’, L2’, and FnIII-1 domains. The membrane-proximal regions of the FnIII-1 domain are oriented towards the L1 domain (Fig. 3D). Consequently, the altered receptor geometry results in a separation of >10 nm between the membrane proximal regions of the FnIII-3 domains. This separation is comparable to their separation in the Λ-shaped apo-IR structure (Fig. 2D,3A). The resulting spatial configuration further retains the TMD dimerization, thereby inhibiting IR-TKD activation.

### Ada and Trim have two α-helical motifs which cross-link IR-A L1 and FnIII-1’ domains

The antagonistic molecules, Ada and Trim, stabilize the L1 and the FnIII-1’ domains in close proximity, inducing a structural rearrangement that shifts the FnIII-1’ domain in the membrane distal direction compared to the insulin-bound IR-ECD and positions it within the plane defined by the L1, CR, and L2 domains (Fig. 3D). Structurally, both Ada and Trim consist of two α-helices connected by a short linker forming a V-shaped molecule (Fig. 4C,D). The linker component, which anchors the peptides via their C-termini, does not directly interact with IR-ECD. Rather, it serves as an anchor that determines the orientation and distance of Site-1 and Site-2’ binding helices. In order to accommodate the spatial arrangement of Site 1 and Site 2’, the membrane-proximal region of the FnIII-1’ domain rotates outward from the IR-ECD center, resulting in a pronounced separation of the FnIII-2 and FnIII-3 domains (Fig. 4A,B). The Ada scaffold possesses a chiral center and only one of the two Ada diastereoisomers aligns with the cryo-EM density (Fig. 4C,S7). This finding indicates that only this specific isomer is capable of binding to IR. Consequently, this stereoselectivity of Ada may contribute to an underestimation of its binding constant (Table S1), as the binding affinities were measured with an inseparable racemic mixture.

The Site-1 binding Ac-FYDWFERQ peptide (individual amino acids are labeled with “S1” in the following text) of both Ada and Trim forms a helix that interacts with the internal side of the L1 domain of IR-ECD. The helix axis is oriented perpendicularly to the parallel β-sheet of the L1 domain, with the aromatic side-chains of residues F1^S1^, W4^S1^, and F5^S1^ extensively engaging the hydrophobic surface of the L1 β-sheet involving the residues L36, L37, L62, F64, F88, L94 and F96 (Fig. 4E). Furthermore, the stability of the interaction is facilitated by hydrogen bonds between R14 of the L1 domain and Q8^S1^ on the antagonist as well as between K121 of the L1 domain and Y2^S1^ of the antagonist. These interactions not only anchor the Ac-FYDWFERQ peptide to the L1 domain but also contribute to the displacement of the α-CT helix. In similar manner, the Site-2 binding helix, designated as the SLEEEWAQ peptide (with “S2” designations appended to the relevant residues in the ensuing text), has been observed to interact with the FnIII-1’ domain. The Site-2 helix is positioned parallel to the antiparallel β-sheet of the FnIII-1’ domain, which is composed of β-strands 1, 2, and 5 (Fig. 4E). The interaction interface contains a hydrophobic core formed by L2^S2^ and W6^S2^, which pack against the hydrophobic residues L486, W551, and L552 on the FnIII-1’ domain. Additional stabilization is provided by the electrostatic interactions between the negatively charged residues E3^S2^ and E5^S2^ on the antagonists and the positively charged residues K484 and R479 or R488 on the FnIII-1’ domain. Overall, the SLEEEWAQ helix engaged Site-2 of IR-ECD in a structural configuration analogous to that observed in the insulin B-chain helix (insulin B-chain residues S9-G20, Fig. S8).

### Antagonist binding under sub-saturating ligand concentrations

In the preliminary cryo-EM experiments, the IR-ECD:antagonist complex was prepared at a molar ration of 1:20 (1.1mM:22mM). A notable observation was that the majority of particles exhibited a solitary FnIII-2/3 pair that extended away from the IR-ECD head toward the membrane plane (Fig. S9). Subsequent refinement of these particles revealed that they correspond to an IR-ECD dimer with one antagonist molecule fully bound to the receptor, while the second molecule was firmly bound to Site-2 with only partial contribution to the densities observed in Site-1 (Fig. S9). We refer to this structure as an “asymmetric complex” in the subsequent discussion. The asymmetric complex was observed to a greater extent in IR-ECD:Ada dataset, where it was the only species identified. No symmetric complex could be identified unambiguously at 1:20 molar ratio. Conversely, in IR-ECD:Trim (1:20) data, 18% of particles corresponded to the ∩-shaped complex, indicating a divergence in binding behavior between the two antagonists. When IR-ECD was mixed with a 100-fold molar excess of the antagonist (1μM IR-ECD dimer:100μM Ada/Trim), particles corresponding to both symmetric and asymmetric complexes were observed. The symmetric complex, described previously, accounted for 63% and 86% of the particles collected from the 100-fold excess antagonist datasets for Ada and Trim, respectively. The asymmetric complex’s structure was refined to a resolution of 3.5−5 Å. The L1, CR, L2, and FnIII-1 domains of both protomers exhibit a comparable organization to that observed in the ∩-shaped structure. However, the FnIII-2/3 domains from a single protomer are resolved, and their density is tilted away from the L1 domain by approximately ∼20° compared to the ∩-shaped structure (Fig. S9). The density for the second FnIII-2/3 is absent from the map, but additional low-resolution density can be seen near the C-terminus of the FnIII-1 domain, oriented towards the L1’ domain (Fig. S9). This low-resolution density suggests a protomer conformation similar to the unliganded Λ-shaped apo-IR structure, where the FnIII-2 domain interacts extensively with the L1’ domain.

### S661 inhibits IR with a similar mechanism to Ada and Trim

The S661 peptide has demonstrated a more effective antagonism of IR activation compared to Ada or Trim, exhibiting an IR binding affinity that is comparable to that of insulin (Table S1). In order to enhance our comprehension of the inhibitory mechanism, we have determined the IR-ECD:S661 complex cryo-EM structure at a resolution of ∼5.3 Å (Fig. 5). The resulting map was interpreted by means of flexible fitting of IR domains from the IR-ECD:Ada structure obtained in this study (Fig. 4A). Two S661 molecules bind asymmetrically to the IR-ECD dimer, similar to the observation made in the IR-ECD:Ada or Trim complexes, but at lower concentrations of the antagonist (see above). Attempts to increase the S661 concentration resulted in sample aggregation on the cryo-EM grids. We hypothesize that the longer linker in S661 may promote binding of the peptide to two distinct IR dimers through Site-1 and Site-2 helices *in vitro*. Consequently, the present analysis concentrated on the asymmetric IR-ECD:S661 complex (Fig. 5). Its structural characteristics exhibit notable similarities with those of the Ada and Trim-bound complexes (Fig. 4, Fig.S9). The L1, CR, L2, and FnIII-1’ domains form a more rigid receptor head and the S661 binding stabilizes a conformation with the FnIII-2/3 domains extending distally from the IR-ECD dimer center. Two helical densities were observed on the L1 and FnIII-1’ domains, connected by a weaker stretch of density at the membrane-proximal site of L1 and FnIII-1’ (Fig. 5). The positioning of the helical densities corresponds to the binding site for Ac-FYDWFERQ and SLEEEWAQ peptides, as previously discussed. However, the longer and flexible linker in S661 results in a distance between the L1 and FnIII-1’ domains that is ∼12 Å larger in the S661-bound IR-ECD complex than in the Ada and Trim-bound complexes. This leads to an even more open receptor conformation and higher flexibility which prevents refinement of the data to higher resolution (Fig. 5B). Analogously to the observations made in Ada/Trim bound structures, no density corresponding to the α-CT helix was detected in the S661-bound structure.

**Figure 5.**
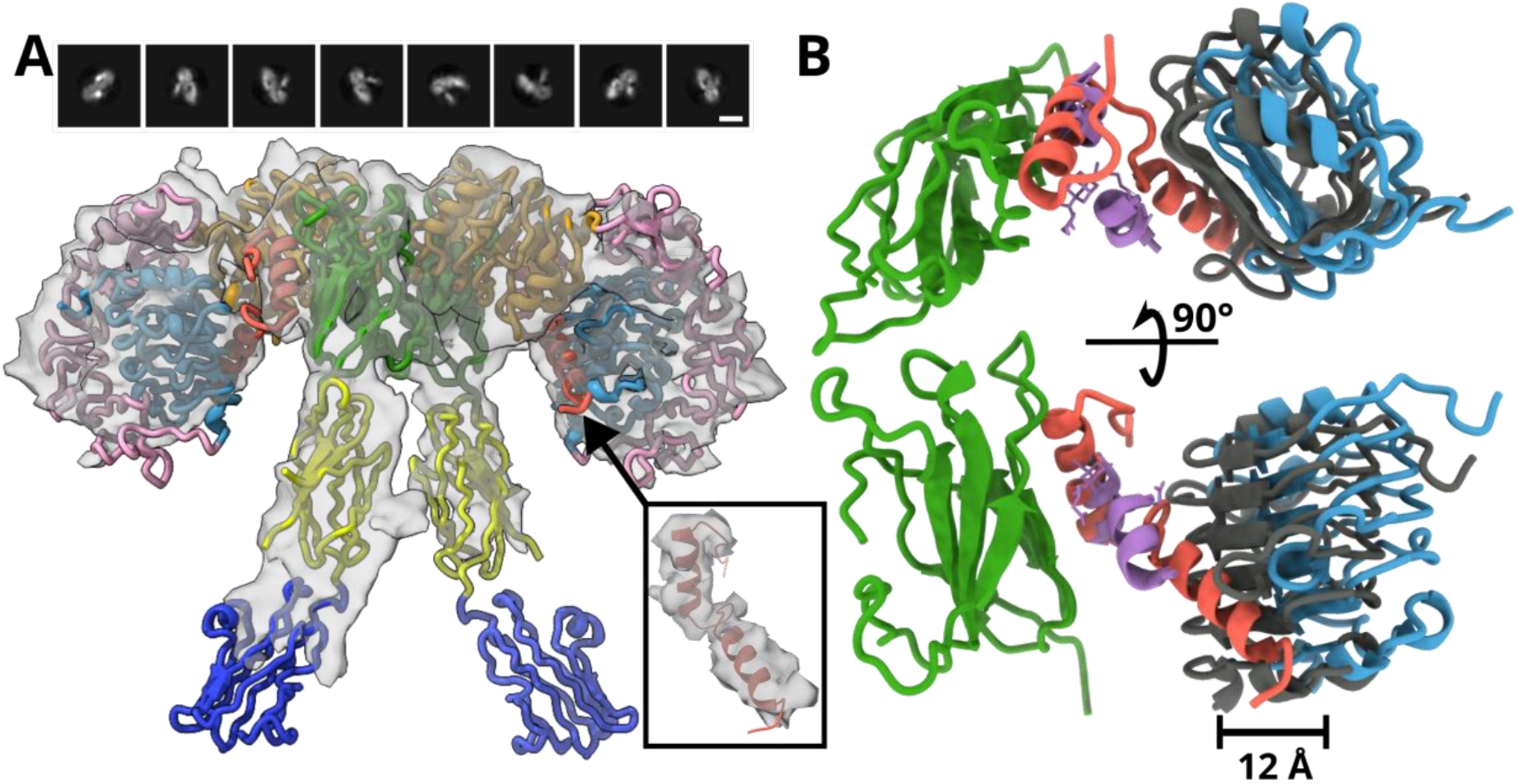
2D class averages and 3D reconstruction with structural model in cartoon representation of IR-ECD in complex with the S661 peptide (A; color coding corresponds to Fig. 2A; S661 is depicted in coral red; domains from different IR protomers are not distinguished). An inset shows a detail of S661 fit into the cryo-EM density. A longer linker between the Site-1 and the Site-2 interacting peptides in S661 (in coral red) compared to Ada (in violet) results in ∼12 Å larger separation of the L1 and FnIII-1’ domains in the case of the IR-ECD:S661 complex structure (B, FnIII-1’ domain of IR-ECD:Ada complex is highlighted in grey).

## Discussion

The vast majority of apo-IR structural studies reported to date (23-25) have identified the Λ-shaped conformation as the initial structural state in the insulin-induced receptor activation pathway. However, our analysis indicates the presence of a ∩-shaped structure in parallel with the previously identified Λ-shaped conformation in the apo-IR sample. Notably, 2D class averages resembling the ∩-shaped structure were observed in a previous cryo-EM study of the human apo-IR (11) and in negative stain EM data of the full-length human IR reconstituted in lipid nanodiscs (21). The binding of Ada, Trim, or S661 molecules with antagonistic effect on the receptor stabilizes the ∩-shaped conformation. This stabilization renders the receptor amenable to structural studies in 3D at resolutions better than 10 Å. We employ 3D reconstructions of the antagonist-bound ∩ state as proxies to facilitate discussion of the structural organization of the alternative apo-IR form(s). The presented data accentuate the transient nature of the L1 domain’s interactions with the FnIII-2’ domain and the α-CT helix in apo-IR, a critical component for receptor activation. The observed high structural flexibility of apo-IR is functionally supported by the observation of IR’s basal signaling activity in the absence of ligand engagement (26).

The existence of the conformational ensemble in apo-IR raises further questions, such as how insulin interacts with these states and whether one particular conformation is favored for insulin-induced IR activation. Previous kinetic studies employing specific insulin analogues with mutations in Site-2 engaging residues provided evidence that the initial contact point of insulin with the receptor is Site-2 (27, 28). Subsequent to this primary interaction, insulin is believed to translocate to Site-1, a binding site formed by the α-CT peptide and the L1 domain, thereby shifting the position of a-CT on L1 and triggering a conformational change in the receptor that results in its activation. This activation model (9, 13) is further supported by “transient” asymmetric structures of insulin:IR complexes, where insulin is located at different positions between Site-2 and Site-1 (14, 15). Recent findings have also implicated R717 of the α-CT as a key residue directly involved in the translocation of insulin from Site-2 to Site-1 (29). Furthermore, studies employing insulin mimetic peptides that bind exclusively to Site-1 or only to Site-2 (13, 15) have confirmed that engagement of both binding sites is imperative for full receptor activation, thereby underscoring the significance of the interplay between Site-1 and Site-2 in this process. In this context, it is noteworthy that when we fitted Ada or Trim to Site-2 in the Λ-shaped apo-IR structure, the antagonist was unable to contact Site-1 due to its short linker. Should Site-2 be confirmed as the initial contact site, there must be specific fluctuations within the receptor apo-structure that would enable the concurrent engagement of Site-1 and displacement of a-CT by the mimetics, as evidenced by the structural data. This proposed mechanism underscores the substantial conformational plasticity of IR in its apo-state.

The Ada and Trim bound IR structures reveal a novel inactive insulin receptor conformation, referred to as the ∩-shaped structure. The ∩-shaped IR structure induced by binding to Ada, Trim, or S661 is distinctly different from previously reported structures of IR engaged by partial antagonists based on insulin dimers (30). In these studies, partial antagonism was attained through stabilization of the Λ-shaped IR structure facilitated by an extensive − though presumably transient - interaction interface between the L1 domain, the FnIII-2’ domain and the α-CT helix on L1. In contrast, the stabilization of the ∩-shaped structure is provided by a distinct structural arrangement involving L1 and FnIII-1’ domains. This structural arrangement is stabilized through close proximity between Site-1 and Site-2 upon Ada, Trim, or S661 binding, preventing receptor activation and inducing structural transition of the IR dimer into the ∩-shaped IR conformation which keeps FnIII-3 domains separated by >10 nm. This structural rearrangement has been shown to protect or alter the interaction interfaces that would typically engage with insulin or insulin-like molecules during receptor activation (Fig. 6). We propose that the binding of Ada, Trim, and S661 might be initiated by the interaction of the antagonists with Site-2. The observed asymmetric structures indicate that the antagonist molecule binds firmly to Site-2, while its interaction with Site-1 remains transient. This is likely due to competition for the L1 binding with the α-CT helix. We hypothesize that the equilibrium between these binding states most probably shifts in a concentration-dependent manner, thereby ultimately driving IR to adopt the ∩-shaped conformation with-both Site-1 sites occupied by the antagonist molecule.

**Figure 6.**
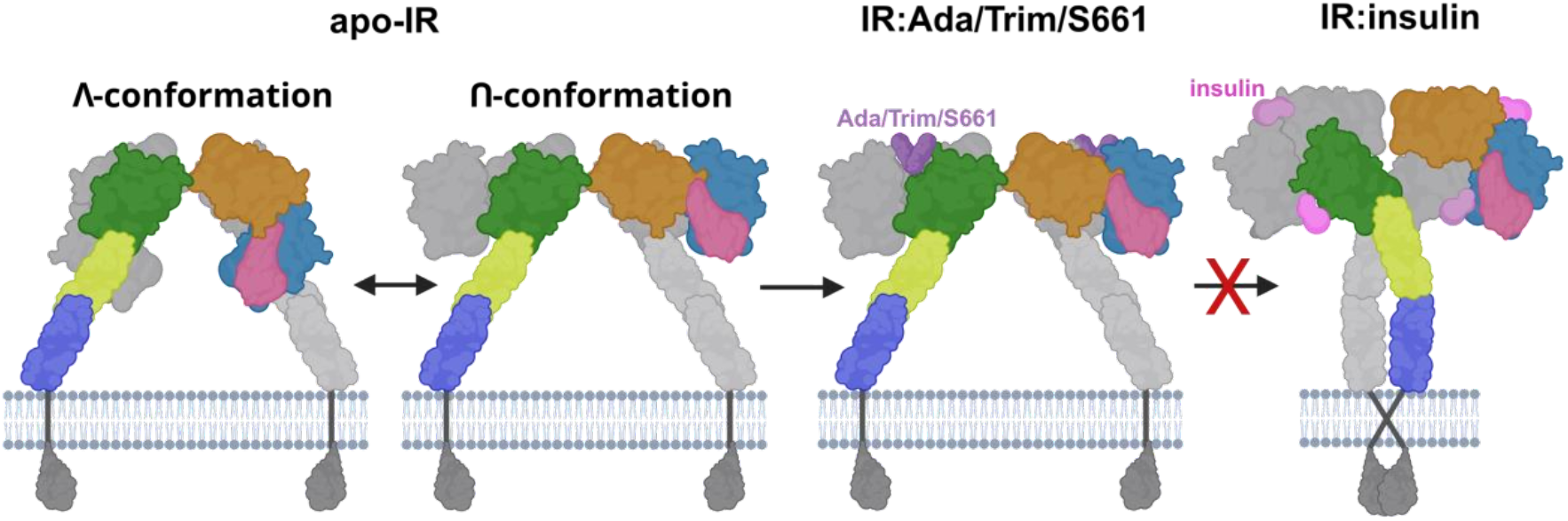
Schematic depiction of the apo-IR conformational plasticity. The Ada, Trim, and S661 bind and stabilize the ∩-shaped IR conformation through conformational selection. The antagonist molecules occupy Site-1 and Site-2 and prevent insulin binding and subsequent receptor activation.

Although Ada, Trim, and S661 stabilize IR in virtually identical structural arrangements, their binding affinities differ significantly. We hypothesize that this discrepancy can be attributed to the distinct structural characteristics exhibited by the antagonists. The short scaffolds in Ada and Trim likely introduce steric restrictions during receptor binding, which likely influences the kon rate. In contrast, flexible peptide linker in S661 reduces conformational barriers, facilitating the appropriate binding conformation required for L1 and FnIII-1’ domain interactions. Importantly, we observed an additional density in the IR-ECD:S661 structure connecting the C-terminus of the SLEEEWAQ peptide to the N-terminal region of the L1 domain. This suggests that the C-terminal residues of S661 that contain a disulfide bridge contribute to an extended interaction surface with the receptor, establishing additional interactions between L1 and FnIII-1’ domains, which results in increased S661 binding affinity compared to Ada and Trim. A number of IR structures in complexes with peptides containing identical amino acid sequence motifs (FYDWFERQ and SLEEEWAQ) have been previously documented (31, 32). The binding modes of these peptides—regardless of whether they engage in two-site binding (e.g., IM459 or S597) or single-site binding (e.g., S519C16; PDB: 5J3H, 33)—were consistent with our data set. A structural comparison of the different peptide sequences reveals that the Ac-FYDWFERQ and SLEEEWAQ motifs employed in our study represent the minimal binding elements necessary to efficiently bring the L1 and FnIII-1 domains in close contact and inhibit IR activation (Fig. S11).

Collectively, the Ada, Trim, and S661 structures reveal general rules for the design of Ac-FYDWFERQ and SLEEEWAQ derived insulin mimetics with either agonistic or antagonistic effect on IR (Fig. S12). Antagonism is elicited through the placement of the L1 and the FnIII-1’ domains in a similar distance from the hypothetical membrane plane. This results in a structural arrangement in which the C-termini of the FnIII-1 domains are oriented in an opposite directions (Fig. S12). Consequently, this leads to the separation of the FnIII-2 and FnIII-3 domains, which prevents the activation of the tyrosine kinase. The proposed IR configuration can be realized through the establishment of a connection between the C-terminus of the Site-1 binding peptide (FYDWFERQ) and the N-terminus of the Site-2 peptide (SLEEEWAQ) by peptide linkers, such as those employed in S661. Alternatively, the C-termini of both peptides can be linked through the use of artificial scaffolds, a method that has been utilized in Ada and Trim. Conversely, agonistic mimetics comprising the Site-2 peptide on the N-terminus and the Site-1 peptide on the C-terminus will yield a vertical separation of the L1 and FnIII-1’ domains. This configuration will enable the C-termini of the FnIII-1 domains to orient towards each other, as evidenced in the previously reported S519C16- and S597-bound complexes (Fig. S12). Consequently, the FnIII-2/3 and the FnIII-2’/3’ domains will be brought into close proximity, a prerequisite for the activation of the kinase.

It is noteworthy that while insulin demonstrates approximately 840-fold greater potency in IR binding compared to Ada (0.47nM vs. 395nM), the administered dose of Ada (125U/kg) in the insulin tolerance test experiment was merely 166-fold higher than the insulin dose (0.75U/kg) (Fig. 1C). Notwithstanding, this amount proved sufficient to markedly attenuate insulin action. This underscores the potential benefits of mimetic antagonists *in vivo* for glycemic control, particularly in view of the reduced relative dose necessary for their effect. Moreover, the synthetic preparation of antagonistic compounds, such as Ada and Trim, may offer certain advantages over insulin or natural peptides with high affinity. These advantages include higher stability in blood plasma and extended circulation time (Probing Tripodal Peptide Scaffolds as Insulin and IGF-1 Receptor Ligands Eur. J. Org. Chem. 2018 2:37).

In conclusion, our results provide a mechanistic understanding for IR inhibition by the Ada, Trim, and S661 antagonists. All three molecules induce a ∩-shaped IR structure in which the membrane proximal parts of the FnIII-3 domains are stabilized in >10 nm distance prohibiting transmembrane signal transduction and kinase domain activation. The FnIII-3 domain separation is independent of the interaction between the L1 and the FnIII-2 domains. Importantly, all three antagonist-bound structures are reminiscent of the ∩-shaped apo-IR conformation, providing further evidence that apo-IR undergoes transitions between different structural states even in the absence of insulin or other activating/inhibitory ligands and allowing for the first time an exploration of high-resolution structural information of the ∩ state (in the antagonist-bound data). In line with recent single molecule studies of the IR-ECD in solution (34), these findings suggest that apo-IR should not be viewed as a single rigid structure, but as a dynamic equilibrium between multiple distinct conformations. This conformational plasticity structurally satisfies two critical functional requirements of the IR’s apo state: (I) the attenuation of transmembrane coupling in the absence of ligands and, at the same time, (II) rapid, large scale structural transitions to the activated states once ligand becomes available.

## Methods

### Protein production

The IR-A-ECD dimer was produced as described earlier (11). In brief, the sequence encoding for IR signal sequence followed by residues 1-917 of human IR-A, linker with 8xHis, protease 3C cleavage site, and a tandem-affinity purification (TAP) tag was cloned into the PTT-6 vector for transient expression in HEK293F cells. The cells from the 2 liters of transfected culture were harvested, lysed, and the supernatant was loaded twice onto the IgG Sepharose beads. Following extensive washing, the beads were incubated overnight with glutathione S-transferase-tagged protease 3C for TAP cleavage. The IR-ECD was collected in a single step elution and loaded on the Ni-NTA beads for a second affinity purification. The IR-ECD was eluted using a buffer containing 280mM imidazol, immediatelly desalted, concentrated, and loaded on the Superdex 200 10/300GL column equilibrated with a HBS buffer (50mM HEPES, pH=7.5, 150mM NaCl). The peak fractions were directly frozen into liquid nitrogen (LN2) and used for cryo-EM later.

### Synthesis of ligands

Ada was prepared according to the protocol described by Hajduch et al. (20). Trim was prepared according to Selicharová et al. (19). S661 was synthesized using a protocol described in Lubos et al. (35). Peptides were prepared by solid-phase peptide synthesis (SPPS) on the Spyder Mark IV Multiple Peptide Synthesizer (EP 36.7), developed in Development Center of the Institute of Organic Chemistry and Biochemistry. Analytical data for Ada, Trim and S661 compounds are provided in the Supporting Information (Fig. S13, S14).

### Receptor-binding studies

Binding affinities of the compounds for IR-A were determined by the competition of hormones with [125I]-monoiodotyrosyl-TyrA14-insulin for IR-A in cell membranes of human IM-9 lymphocytes (ATCC) as described previously (37). Radiolabeled [125I]-monoiodotyrosyl-TyrA14-insulin was prepared according to a procedure described in detail by Asai et al. (38). The binding curve of each compound was determined in duplicate, and the final dissociation constant (*K*_d_) was calculated from at least three binding curves. Binding data were analyzed in GraphPad Prism 8.0 using non-linear regression for one site binding and taking into account potential ligand depletion. In parallel, binding data were also analyzed in Excel software by a one-site fitting program developed in the laboratory of Dr. Pierre De Meyts (A. V. Groth and R. M. Shymko, Hagedorn Research Institute, Denmark, a kind gift of Pierre De Meyts). Scatchard plots were drawn using the software.

### Receptor phosphorylation assay and antagonism assay

Mouse embryonic fibroblasts (IR-A) derived from IGF-1R knockout mice and stably transfected with human IR-A, kindly provided by A. Belfiore (Catanzaro, Italy), were grown as described previously (39). Ligand-dose response IR-A autophosphorylation levels for the analogs were determined using an In-Cell Western assay adapted for chemiluminiscence as described in Machackova et al. (40) and the inhibition of insulin (at 10 nM) activity by the compounds (concentration range from 0.01 nM to 50 μM) was performed as described in Lubos et al. (35). Autophosphorylation of the receptor was monitored using anti-phospho-IGF-1Rβ (Tyr1135/1136)/IRβ (Tyr1150/1151) antibody (Cell Signaling Technology). Data were subtracted from the background values and expressed as the contribution of phosphorylation relative to the 10 nM insulin signal. Stimulation with insulin and antagonism curves were measured in duplicate, and the experiment was repeated 3 times. Insulin stimulation was determined for 0.01-500 nM. Antagonism of Ada and Trim was measured for 1nM – 50 μM and antagonism of S661 for 0.1 nM – 10 μM. The specific concentration points slightly differed among the plates. The Ada, Trim and S661 did not induce autophosphorylation of IR. The curves of the compound were measured in mono-plicate and the experiment was repeated twice. Control wells and wells stimulated with 10 nM insulin were conducted at least as tetraplicates on each plate. Nonlinear regression curve fitting the combined data from all experiments (i.e., 6 values per point) was carried out with GraphPad Prism 5 software.

Control western blots were performed as in KříŽková et al. (41). The IR-A cells on 24-well plates were stimulated with 50 - 1 μM concentrations of the ligands in the presence of 10 nM insulin for 10 min. Proteins were routinely analyzed using immunoblotting. The membranes were probed with anti-phospho-IGF-1Rβ (Tyr1135/1136)/IRβ (Tyr1150/1151) (Cell Signaling Technology). Anti-actin (20-33) antibody (Sigma–Aldrich, cat. A5060) was used as a loading control.

### Insulin tolerance test (ITT)

Animal procedures followed the European Communities Council Directive 86/609/EEC and were approved by the Institutional Ethical Committee on Animal Experimentation, Czech Academy of Sciences, Prague (protocol No.121-2023). The experiment was done as described in Páníková et al. (42). Briefly, male, 12-weeks old C57BL/6 mice (weighing 23–29 g) purchased from Charles River were randomly divided into three groups of ten mice each. Prior to the test, the mice were fasted overnight (18 hours). Groups of mice were injected with saline, human insulin (0.75 U/kg) or with human insulin (0.75 U/kg) in combination with Ada (125 U/kg). One Unit (U) is defined as 6 nmol of insulin or Ada. Blood glucose was measured with a glucometer (Arkray, Kyoto, Japan) in a drop of blood obtained from the tail vein at times 0, 10, 20, 30, 45, 60, 120 and 150 min after injection of compounds or saline. The data were analyzed in GraphPad Prism 8.0 (San Diego, USA). The significance of the changes induced by treatment was calculated using two-tailed t-test for independent samples.

### Cryo-EM sample preparation and data acquisition

The IR fractions from the gel filtration were thawed on ice. The 1.1 µM IR-dimer sample was incubated with 200 mM or 1 mM Ada or Trim sample in 9:1 ratio to form 1:20 or 1:100 (IR dimer: ligand) complex, respectively. For the S661 ligand, the 0.4 µM IR-dimer was mixes with 360 µM S661 in 9:1 ratio (1:100 IR-dimer:S661 complex) and vitrified within 1 minute as longer incubation cause observation of large aggregates in the sample and low number of monomeric IR particles in the ice. The R1.2/1.3 TEM grids (Quantifoil, Cu, 300 mesh) were activated in H/O plasma (Gatan Solarus 2) for 1 minute. The overall volume of 3.4 ml of the sample was applied to the freshly plasma-cleaned TEM grid, followed by 30 s incubation, and 4 s blotting before its vitrification into the liquid ethane. The whole vitrification process was carried out using ThermoScientific Vitrobot IV. The Vitrobot chamber was maintained at 6°C and 90 – 100% relative humidity. Subsequently, samples were loaded into Talos Arctica transmission electron microscope for grid screening.

Single particle cryo-EM data were collected in automated manner on Titan Krios G1 (ThermoScientific) using SerialEM (43) software (an example micrographs is shown at Fig. S3). The microscope was aligned for fringe-free imaging and equipped with Bioquantum K3 (Ametek) direct electron detector. The camera was operated in electron counting mode and the data for IR in complex with Ada, Trim, and apo-IR were collected at the pixel size of 0.834 Å/px, whereas the data for S661 were collected in correlated double sampling (CDS) mode and a pixel size of 0.51 Å/px. The microscope condenser system was set to provide 21 e−/Å2 s (Ada, Trim, apo-IR data) resp. 29 e−/Å2 s (S661) electron flux on the specimen. The data from 2.0 s exposure were stored into 40 frames. The energy selecting slit was set to 10 eV. The data from 3 × 3 neighboring holes were collected using beam/image shifting while compensating for the additional coma aberration induced by beam shift. The data were collected with the nominal defocus range of −1.2 to −2.4 µm. Overall, the movie sets used for data analysis comprised 9.632 movies for IR-ECD:Ada complex, 19.332 movies for IR-ECD:Trim complex, 7.339 movies for IR-ECD:S661 complex, and 17.266 movies for apo-IR-ECD dataset.

### Cryo-EM data processing

Processing was performed in CryoSparc v.4.1.0 (22) and Relion v.5.0 (44). Movies were first imported into CryoSparc, motion correction was performed using Patch motion correction job and contrast transfer function parameters were determined using Ctffind4 (45). Micrographs, which acquired drift larger than 50 pixels, astigmatism above 800 Å, CTF fit to a resolution worse than 6 Å, contained crystalline ice or large ice crystal contamination, were excluded from further processing. A subset of 500 micrographs were subjected to initial particle picking using a Blob picker (110-170 Å particle diameter) job. The 2D classes from the reference-free 2D classification of these particles which corresponded to IR were used as templates for subsequent template picking of the whole dataset. Particles were extracted from the micrographs, binned four times and subjected to multiple rounds of reference-free 2D classification to remove false positive picks and corrupted particles. The initial round of 3D classification was performed through the *ab initio* routine in CryoSparc. Particles corresponding to IR in various conformations were re-extracted without binning (or 2x binning for S661 complex) and iteratively refined using Homogeneous refinement and Heterogeneous refinement jobs. C2 symmetry was imposed during the reconstruction step for apo-IR-ECD and IR-ECD:Ada, IR-ECD:Trim, IR-ECD:S661 with ligands symmetrically bound to each both IR monomers. Other maps were refined without imposing any symmetry. Subsequently, masks were created to comprise different regions of the complex (Table S2) followed by further iterative refinement using Local refinement, Local CTF refinement jobs. Additionally, refinement of beam tilt, trefoil, and magnification anisotropy was included to the final refinement iterations for Ada/Trim symmetric dimer data. The maps were exported to Relion for postprocessing and local resolution calculation. The final resolution for each 3D reconstruction in terms of Fourier shell correlation is summarized Table S2. In parallel to postprocessing in Relion, maps were sharpened using deepEMhancer for the purpose of figure preparation. Complexes with symmetrically bound ligands were subjected to 3D variability analysis in CryoSparc with a focus mask on FnIII-2/3 and FnIII-2’/3’ regions, respectively.

### Cryo-EM data interpretation

Coordinates of the PDB entry 6SOF were used for the model building into maps with symmetrically and asymmetrically bound ligands. The PDB entry 8DTM was used for interpretation of apo-IR-ECD map. Regions of the symmetric Ada/Trim structures corresponding to L1, CR, L2, FnIII-1 domains were interpreted as follows: first, individual domains were docked as rigid bodies into the map using Chimera (46) and Phenix (47) followed by manual building of the missing residues in Coot (48). Finally, the structure was iteratively refined using real space refinement in Phenix and semi-manual adjustments in Coot. Model geometry was validated using Molprobity (49). The peptide (SLEEEWAQ, FYDFWERQ) components of the ligand molecules were first built into the cryo-EM map in Coot followed by refinement in Phenix. Adamantane- and inosine-derived linkers were first built in Lidia, then docked into the cryo-EM map, and finally linked to the peptide sequences using Acedrgn (50). Coordinates for Fn-III-2 domain was docked as a rigid body into the cryo-EM density and refined using molecular dynamics flexible fitting in Isolde (51) with the distance restraints within the FnIII-2 domain. Similarly, asymmetric IR-ECD:Ada/Trim/S661 structures as well as symmetric IR-ECD:S661 structure and apo-IR-ECD structure were interpreted by rigid body docking of individual domains into the cryo-EM maps followed by molecular dynamics flexible fitting refinement using Isolde while retaining distances within individual domains. The SLEEEWAQ:FnIII-1’ and FYDFWERQ:L1 models from the IR-ECD:Ada cryo-EM map were rigid body docked into the IR:S661 and the backbone of the peptide-linker was subsequently built in Coot.

## Acknowledgemets

This work was supported by the project National Institute for Research of Metabolic and Cardiovascular Diseases (Program EXCELES, ID Project No. LX22NPO5104, Funded by the European Union-Next Generation EU), and by the Academy of Sciences of the Czech Republic (Research Project RVO:52, support to the Institute of Organic Chemistry and Biochemistry) and by the Federal Ministry of Education and Research (BMBF) and the German Center for Diabetes Research (DZD e.V.). We acknowledge Cryo-electron microscopy and tomography core facility CEITEC MU of CIISB, Instruct-CZ Centre, supported by MEYS CR (LM53) and European Regional Development Fund-Project "Innovation of Czech Infrastructure for Integrative Structural Biology” (No. CZ.02.01.01/00/23_015/54).

## Author contributions

MPo: investigation, manuscript preparation; MPi: investigation IS: methodology, data curation, writing; MSGH, BF, and ML: synthetic methodology; LZ: biological experiments, data curation; MG: preparation of human insulin receptor ECD; IBS: investigation, manuscript preparation, editing; UC: investigation, manuscript preparation, editing JJ: conceptualization, supervision, data curation, writing. JN: methodology, supervision, investigation, manuscript preparation, editing, review, writing.

## Data availability

The cryo-EM density map and atomic coordinates for IR-ECD:Ada, IR-ECD:Trim, IR-ECD:S661, and Λ-shaped apo-IR have been deposited in the Electron Microscopy Data Bank and the PDB, under accession codes EMD-54026, EMD-54028, EMD-54063, EMD-54066, EMD-54023, EMD-54015 and 9RKY, 9RL2, 9RMJ, 9RMT, 9RKW, 9RKD, respectively.

