## Supplementary Information for "Antagonistic insulin mimetics lock the insulin receptor in an alternative apo-state"

Martin Polák[1], Irena Selicharová[2], Matyáš Pinkas[1], María Soledad Garre Hernández[2], Benjamin Fabre[2], Marta Lubos[2], Lenka Žáková[2], Michal Grzybek[3], Ingmar B. Schäfer[3,4,5], Ünal Coskun[3,4,5,6], Jiří Jiráček[2]*, and Jiří Nováček[1]*

**Table S1**: Overview of insulin mimetic compounds discussed. Peptide sequences with a cyan background bind to Site-2 in the IR and sequences with a grey background bind to IR Site-1. The Cys residues forming the disulfide bridge are highlighted in red. Underlined peptide sequences were used for the construction of compounds Ada and Trim (Fig. 1). Superscripts N and C indicate N or C termini of peptides. Ac is N-terminal acetyl. Asterisk (*) indicates that Ada is a mixture of two diastereoisomers. (A) indicates that the compound activates IR, and (I) indicates that the compound inhibits IR. Receptor binding affinities for IR-A and biological activities were determined by different teams and by different methodologies as indicated in the subnotes[a-f].

| **Code** | **Sequence** | **Binding affinity for IR-A** (nM) | **Biological activity** (nM) |
|---|---|---|---|
| **HI** (A) | Human insulin | 0.32 ± 0.06 (4)[a] | 2.77[b] |
| **S597** (A) | Ac-[N]SLEEEWAQIECEVYGRGCPSESFYDWFERQL[C]-amide | 2.5[c] | 58 % of insulin[d] |
| **S519** (A) | [N]SLEEEWAQVECEVYGRGCPSGSLDESFYDWFERQLG[C] | - | 4.2[e] |
| **S661** (I) | [N]GSLDESFYDWFERQLGGGSGGSSLEEEWAQIQCEVW-GRGCPSY[C]-amide | 2.62 ± 0.4 (3)[a] | 3.26[f] |

| | | | |
|---|---|---|---|
| **Ada** (I) | $^{N}$SLEEEWAQ$^{C}$-adamatane* scaffold-$^{C}$QREFWDYF$^{N}$-Ac | 395 ± 27 (3)[a] | 950[f] |
| **Trim** (I) | $^{N}$SLEEEWAQ$^{C}$-trimesic acid scaffold- $^{C}$QREFWDYF$^{N}$-Ac | 1257 ± 244 (3)[a] | 1331[f] |

[a]$K_d$ ± S.D., from this work (IR-A in IM-9 lymphocytes, details are in Methods). [b]EC50 with 95% IC likelihood 1.89-2.93, from this work (stimulation of IR-A autophosphorylation in fibroblasts, details are in Methods). [c]Binding affinity (IC50) for solubilized IR (from Jensen et al. 55). [d]EC50 for stimulation of IR-A autophosphorylation in rat myoblast cells stably transfected with the human IR-A (from Jensen et al. 55). [e]EC50 determined by lipogenesis stimulation assay in adipocytes (from Schaffer et al. 16). [f]IC50 with 95% IC likelihood 2.82-5.42 for S661, 689-1124 for Ada, and 790-2582 for Trim, from this work (stimulation of IR-A autophosphorylation in fibroblasts, details are in Methods).

**Table S2: Data acquisition, processing, and structure refinement details.**

| | **IR:ada** | **IR:trim** | **IR:s661** | **IR-apo** |
|---|---|---|---|---|
| **Cryo-EM data acquisition** | | | | |
| Microscope | Titan Krios G1 | | | |
| Voltage [kV] | 300 | | | |
| Camera | Bioquantum K3 | | | |
| Magnification | 59.981x | 59.981x | 97.790x | 59.981x |
| Pixel size [Å] | 0.8336 | 0.8336 | 0.5113 | 0.8336 |
| Defocus range [µm] | -1.2 - -2.4 | | | |
| Exposure time [s] | 2 | | | |
| Movies | 9.632 | 19.332 | 7.339 | 17.266 |
| Number of frames per movie | 40 | | | |
| Total dose per movie [e/Å$^2$] | 43.2 | 43.2 | 57.4 | 43.2 |

| **Cryo-EM data processing** | | | | | | |
|---|---|---|---|---|---|---|
| Map | Saturated IR:ada | Unsaturated IR:ada | Saturated IR:trim | Unsaturated IR:trim | Unsaturated IR:s661 | IR-apo |
| Particles | 334.251 | 200.302 | 525.879 | 82.943 | 70.993 | 34.347 |
| Imposed symmetry | C2 | C1 | C2 | C1 | C1 | C2 |
| Map Resolution $FSC_{0.5}/FSC_{0.143}$ | 3.4/3.0 | 4.1/3.5 | 3.6/3.2 | 7.1/4.2 | 8.0/5.3 | 7.7/6.0 |
| B factor [Å$^2$] | -42 | -46 | -71 | -51 | -156 | -481 |
| EMDB | 54026 | 54028 | 54063 | 54066 | 54023 | 54015 |
| PDB | 9RKY | 9RL2 | 9RMJ | 9RMT | 9RKW | 9RKD |
| RMS deviations: bond length [Å] / bond angles [°] | 0.003/0.506 | | 0.003/0.477 | | | |
| Ramachandran plot [%]: favoured / allowed / disallowed | 95.52/4.48/0 | | 94.36/5.64/0 | | | |
| Validation: Molprobity score / clash score / poor rotamers [%] | 6.55/2.21/0.23 | | 7.02/2.3/0.54 | | | |

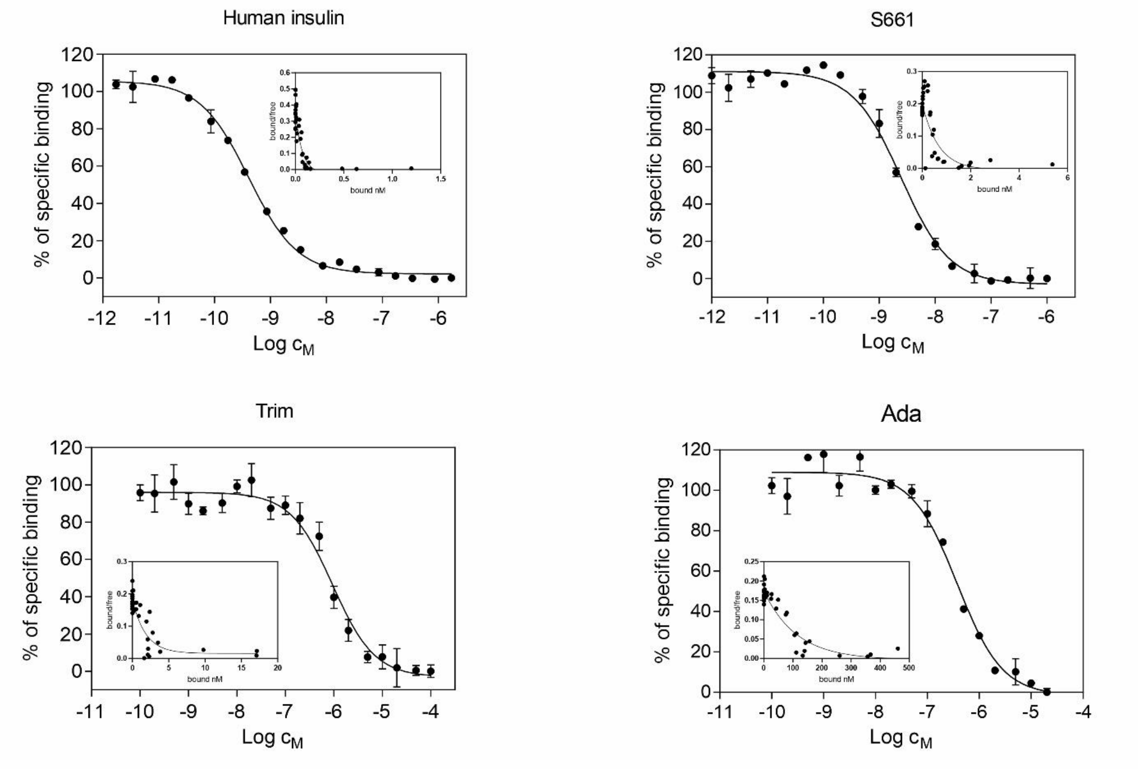

**Figure S1.** Representative binding curves of human insulin and insulin mimetics on IR-A with Scatchard plots (Scatchard plots are in the inserts).

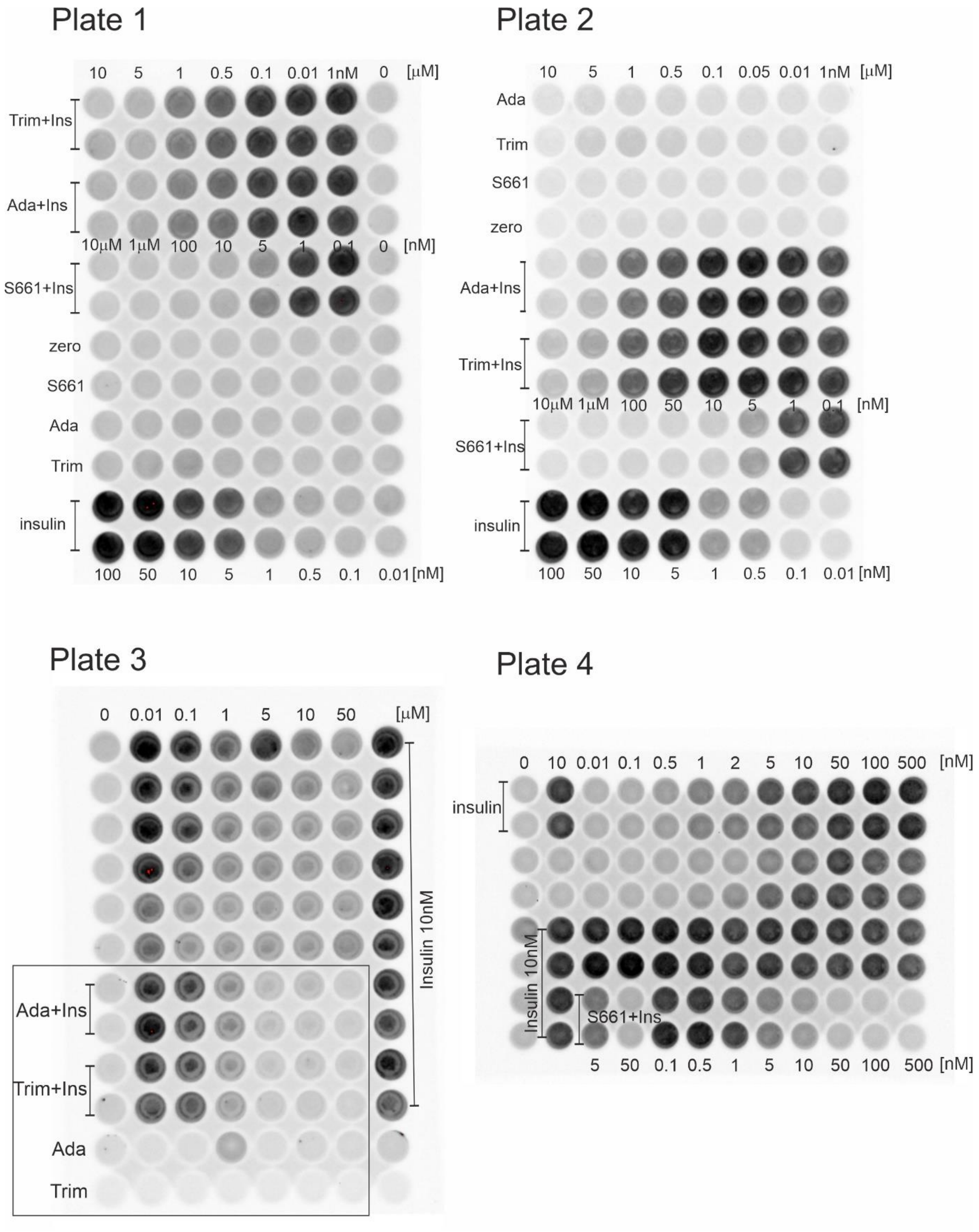

**Figure S2:** "In Cell western" analysis of IR inhibition by Ada, Trim and S661. The cells were treated with compounds at specified concentrations. The concentration ranges slightly differed between the plates. For agonism, compounds were measured in monoplicates, for insulin and antagonism in duplicates. For antagonism, 10 nM insulin was added together with the compounds.

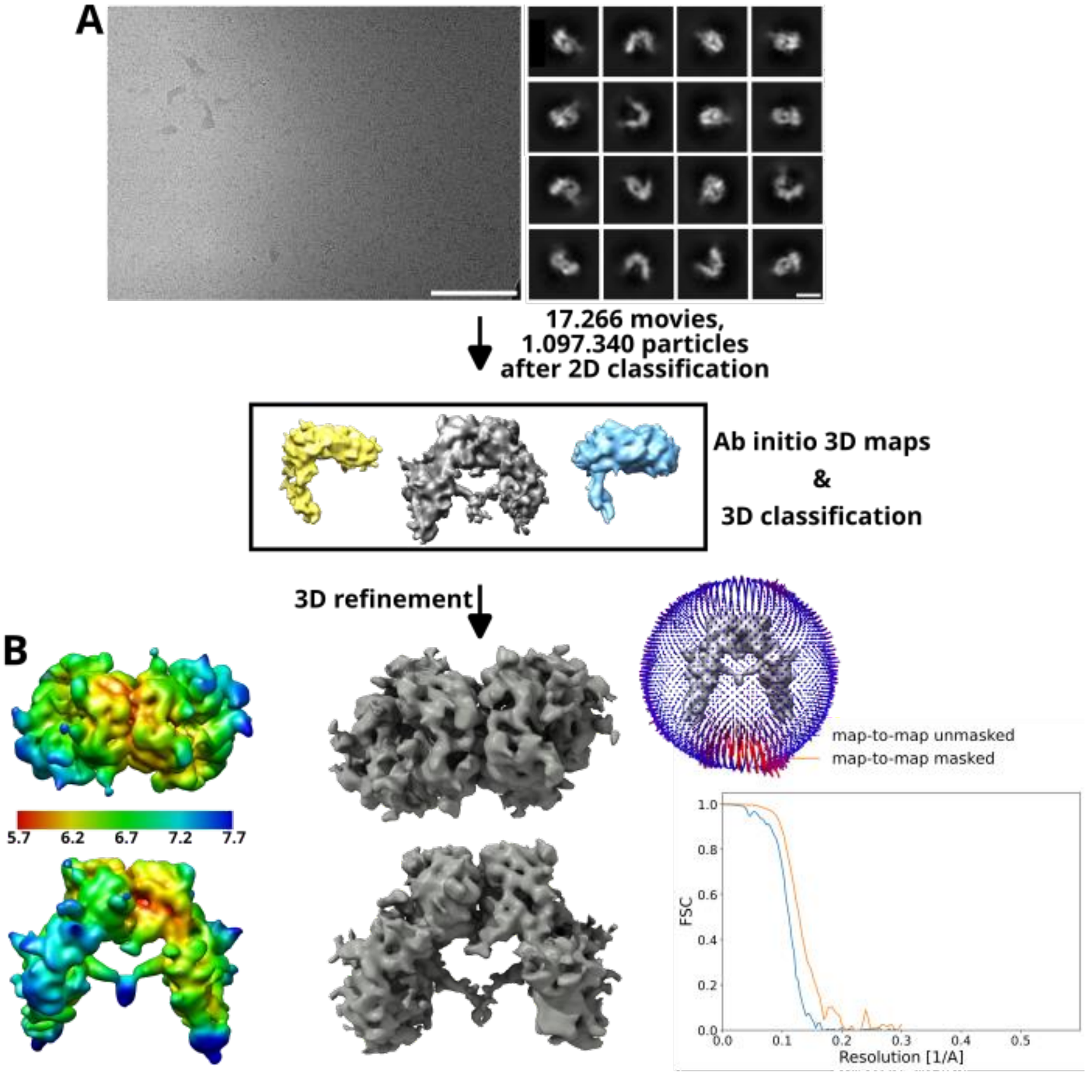

**Figure S3:** Cryo-EM data processing scheme utilized to analyze apo-IR-ECD data (A; scale bars correspond to 100nm for the micrograph and 10 nm for 2D class averages). Final Λ-structure IR-ECD cryo-EM density, local resolution map, FSC plots, and particle orientation distribution are shown (B).

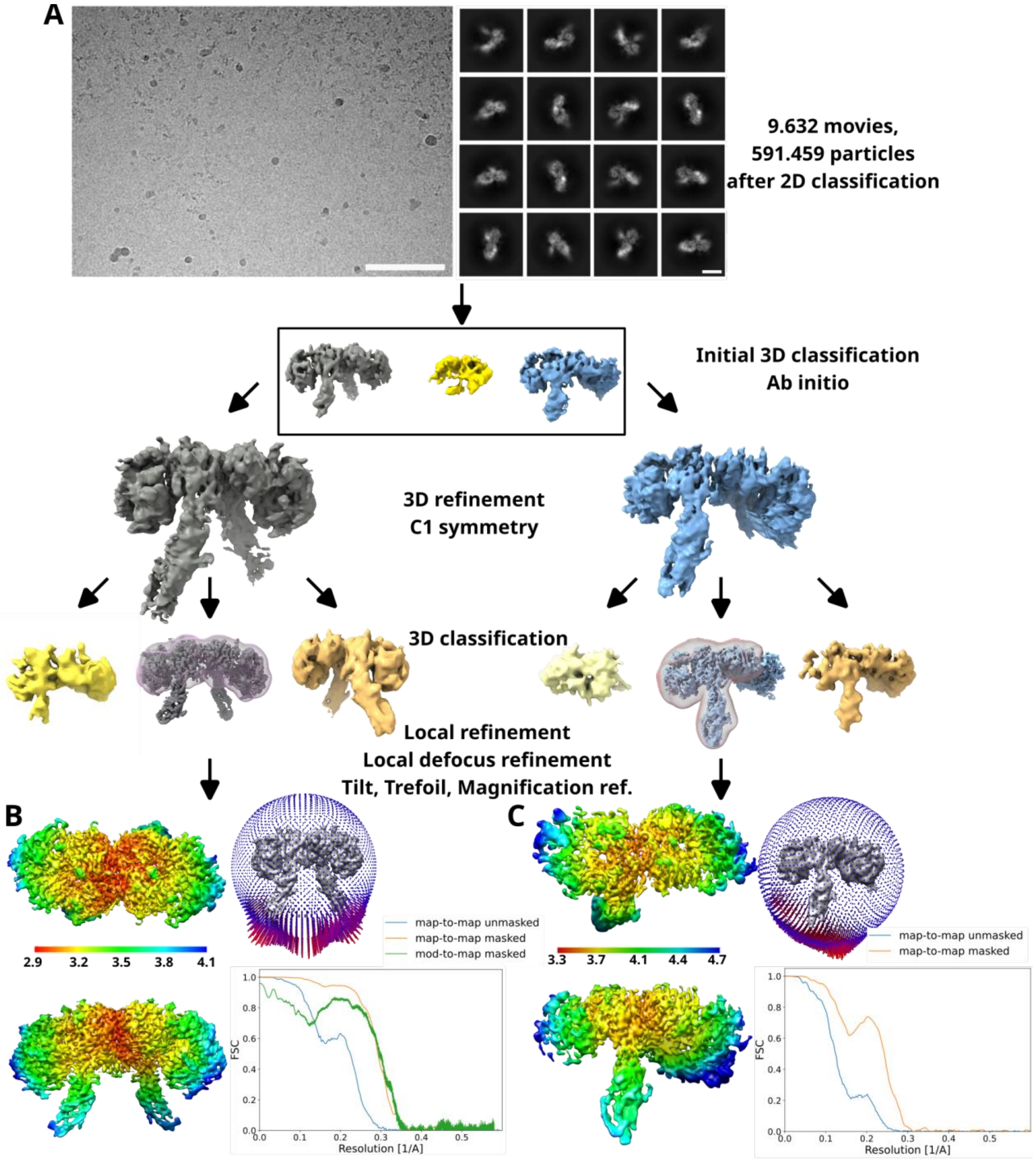

**Figure S4:** Cryo-EM data processing scheme utilized for structure determination of IR complexes presented here (A). Scale bars correspond to 100nm (micrograph) and 10nm (class averages), respectively. Local resolution map of IR-ECD:Ada complex with symmetrically bound antagonist, Fourier shell correlation (FSC) curves of the final model, and particle orientation distribution (B). Local resolution maps of IR-ECD:Ada structure with asymmetrically bound antagonist molecules, corresponding FSC plots, and particle orientation distribution heat map.

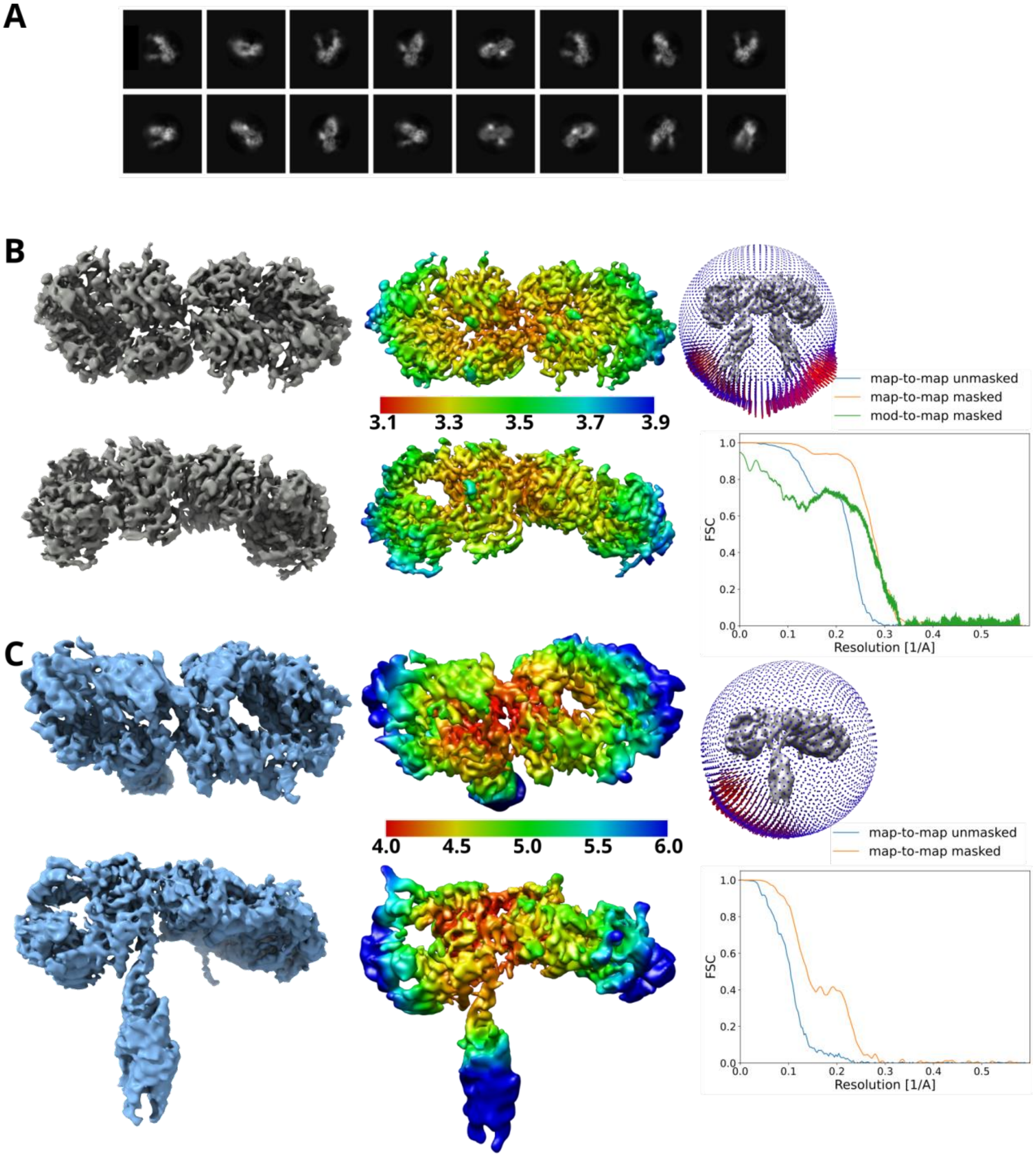

**Figure S5:** Representative classes from the reference-free 2D classification of IR-ECD:Trim complex (A). Local resolution map for the IR-ECD:Trim complex with symmetrically bound antagonist molecules, FSC plots, and particle orientation distribution (B). Local resolution map for IR-ECD:Trim complex with asymmetrically bound antagonist molecules, FSC plots, and particle orientation distribution heat-map. The data processing workflow was the same as shown in Fig. S4.

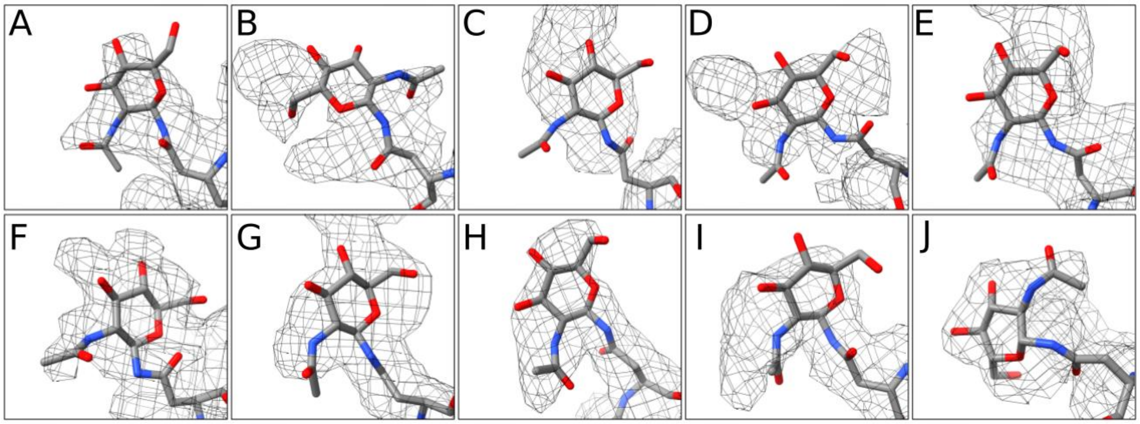

**Figure S6**: IR undergoes significant glycosylation *in vivo*. Overall, fifteen potential N-linked glycosylation sites were identified in α chain and additional four in the β chain (56). We have observed 10 densities extending at the position of the asparagine residues, namely N16 (A), N25 (B), N111 (C), N215 (D), N255 (E), N295 (F), N337 (G), N397 (H), N418 (I), N514 (J). On the other hand, no additional densities were observed at the residues N78 and N282, which were earlier reported to undergo glycosylation. The N16 is in proximity of the antagonist binding site but is not involved in any interaction with Ada or Trim. On the contrary, N111 (L1) and N215 (CR) are positioned close in space between the L1 and CR domains in the antagonist bound IR structures with glycan densities oriented alongside the interface.

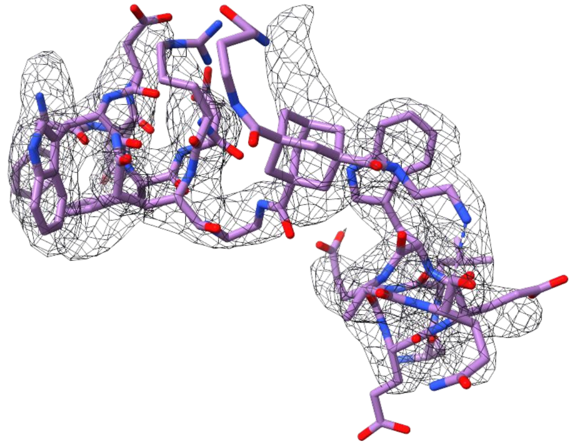

**Figure S7:** A detail of an alternative diastereoisomer of adamantane-derived linker in Ada superimposed into cryo-EM density map.

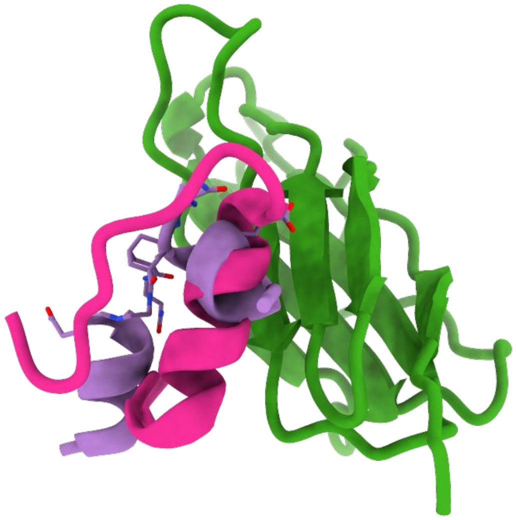

**Figure S8:** Ada (purple) Site-2 peptide (NSLEEEWAQC) binding site on FnIII-1 domain (green) coincides with the binding site of insulin B-chain helix (pink). Superscripts N and C indicate N or C termini of peptide.

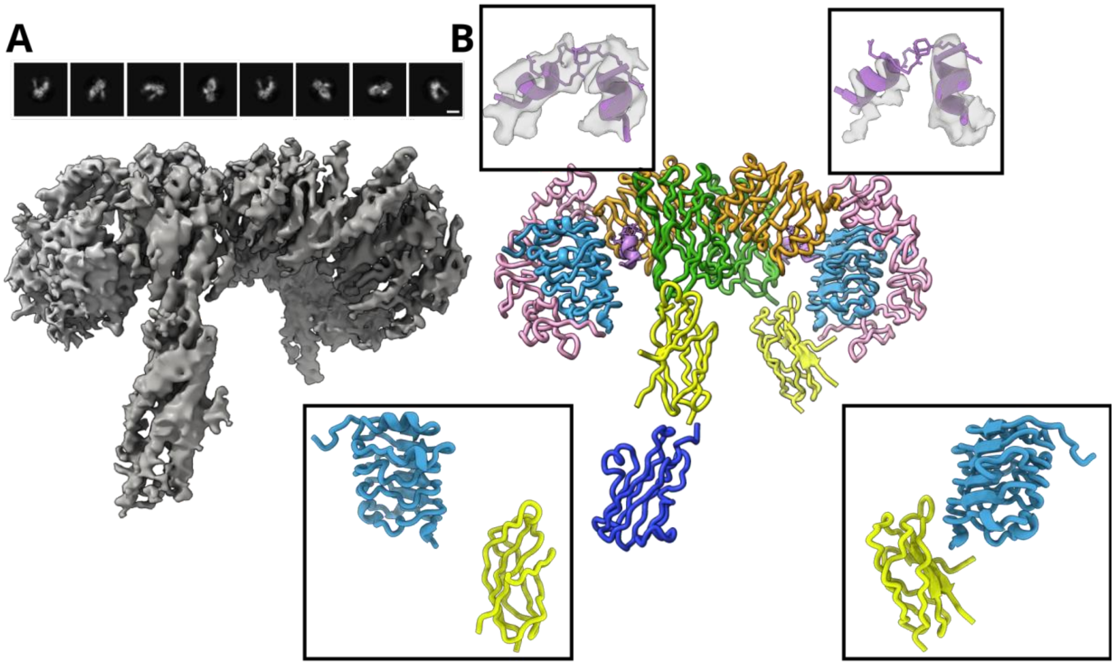

**Figure S9:** 2D class-averages and the cryo-EM map of the "asymmetric" IR-ECD:Ada complex (A; domains from different IR protomers are not distinguished). The map reconstructed from the data collected at the unsaturated ligand concentrations (1:20 molar concentration) shows only one distinct density for the FnIII-2/3 domain. The cryo-EM map was interpreted by flexible fitting of the model determined for IR-ECD:Ada at ligand saturated conditions (B). The insets at the top show a uniform density for Ada in the first binding site (top left) and only partial density for the Site-1 helix observed for the second Ada binding site (top left). The complete Ada binding results in separation of L1 and FnIII-2 domains (bottom left) whereas these domains are kept in similar arrangement as in Λ-shaped apo-IR in the case of incomplete ligand binding (bottom right).

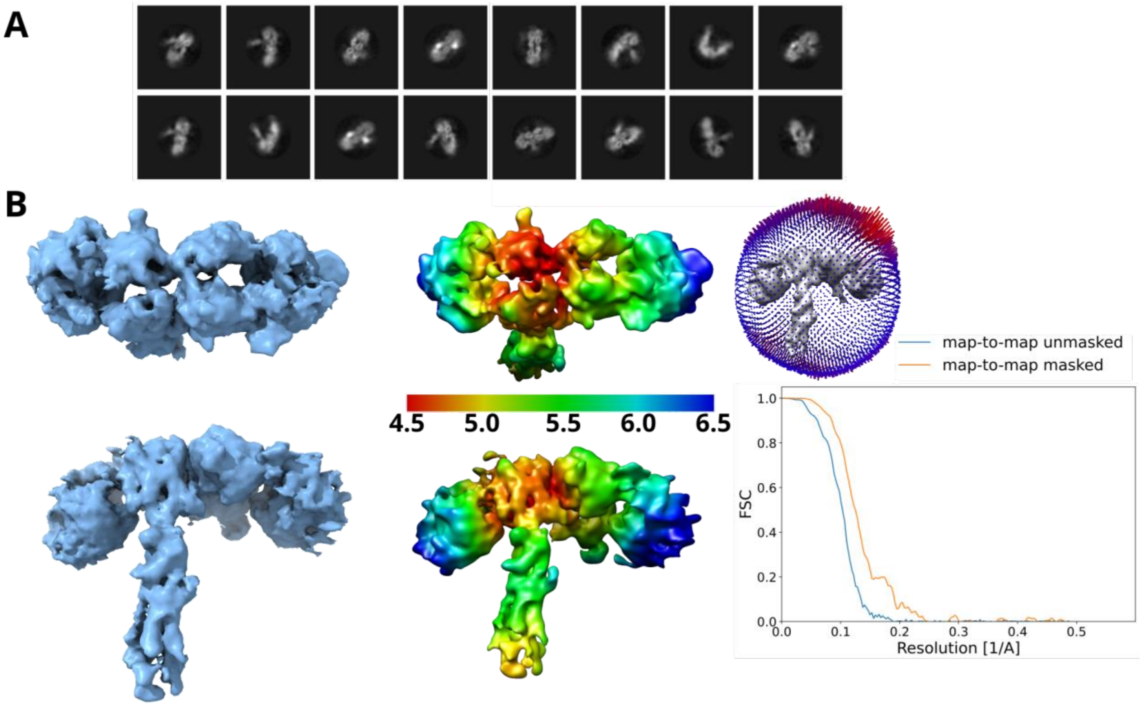

**Figure S10:** Representative classes from the reference-free 2D classification of IR-ECD:S661 complex (A). Local resolution map for the IR-ECD:S661 complex with symmetrically bound S661 molecules, FSC plots, and particle orientation distribution (B). Local resolution map for IR-ECD:S661 complex with asymmetrically bound molecules, FSC plots, and particle orientation distribution heat-map. The data processing workflow was the same as shown in Fig. S4.

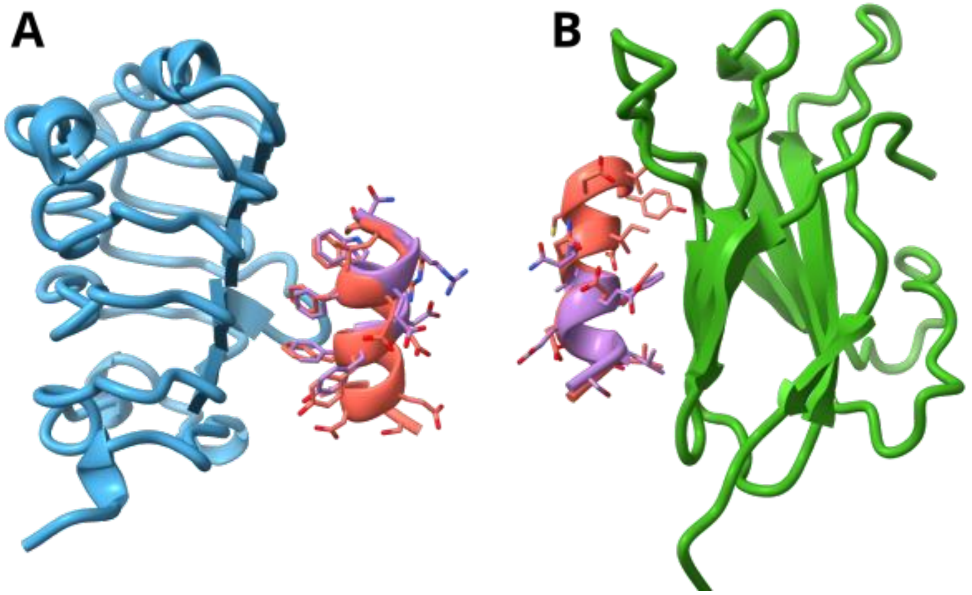

**Figure S11:** A detail of the Ada Site-1 helix (violet) interaction with the L1 (blue) domain and its comparison with the binding mechanism of S519-C16 peptide (A, PDB: 5J3H, coral red). The Ada Site-2 helix (violet) binding to the FnIII-1 domain (green) and its comparison with S597 peptide (B, PDB: 8DTL, coral red).

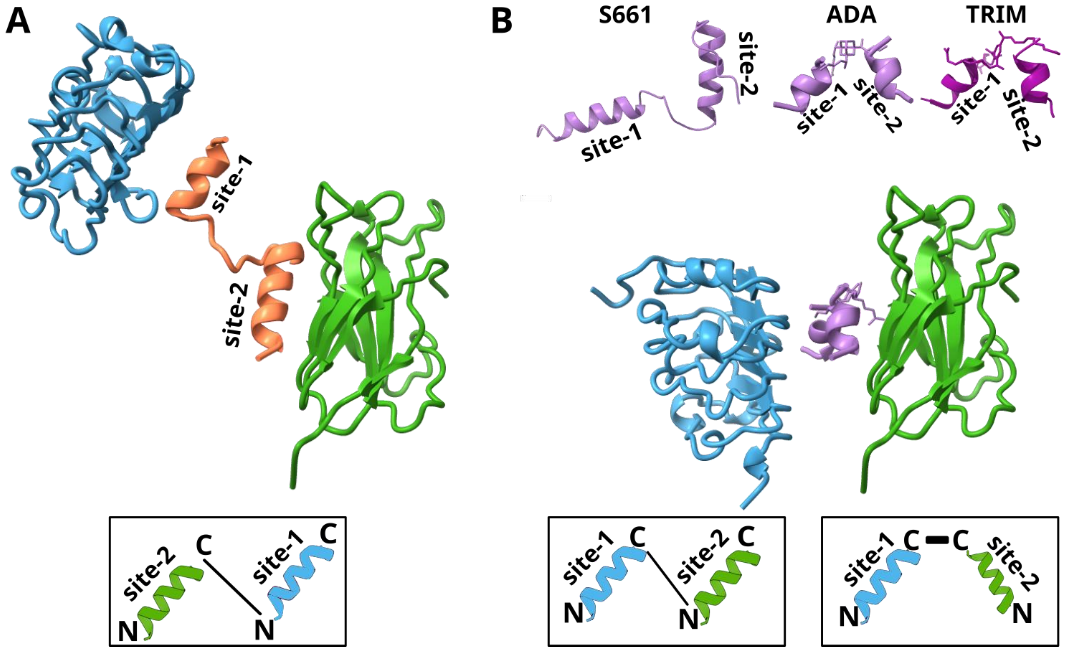

**Figure S12:** Structural arrangement of L1 (blue) and FnIII-1 (green) domains bound with peptides with agonistic (A) or antagonistic (B) effect on IR. Binding of S597 peptide (A, coral red) promotes the transphophorylation reaction, whereas Ada, Trim, or S661 binding inhibits the receptor activity. The insets at the bottom of the panels A and B show general concept for developing peptides by combining Site-1 (light blue) and Site-2 (green) helices which will cause agonism (A) or antagonism (B).

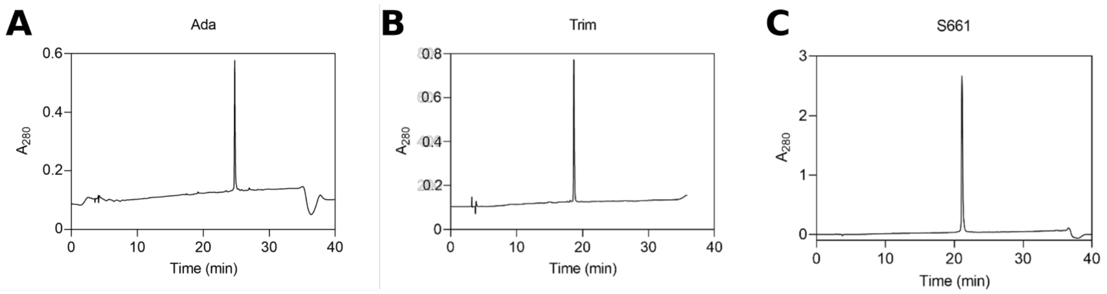

**Figure S13:** HPLC traces of Ada, Trim and S661 compounds.

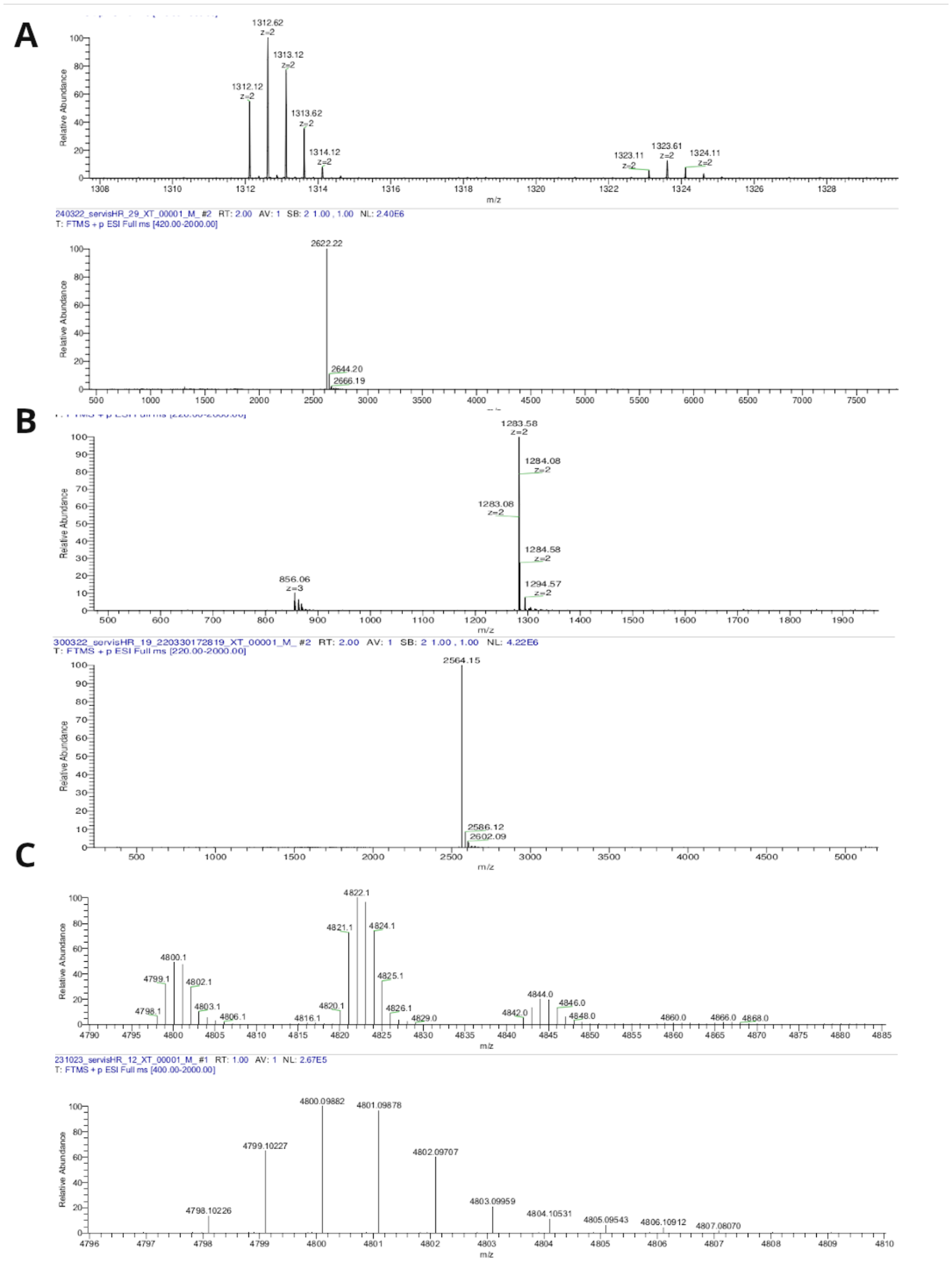

**Figure S14:** Deconvoluted high-resolution mass spectrum of Ada (A; $C_{124}H_{167}N_{29}O_{35}$, Exact Mass: 2622.2179, Molecular Weight: 2623.8680). Deconvoluted high-resolution mass spectrum of Trim (B; $C_{120}H_{157}N_{29}O_{35}$, Exact Mass: 2564.1397, Molecular Weight: 2565.7440). Deconvoluted high-resolution mass spectrum of S661 (C; $C_{211}H_{296}N_{56}O_{70}S_2$, Exact Mass: 4798.0765, Molecular Weight: 4801.1310).
